# In Vitro Wound Simulation: A High-Throughput Device for Scratch Assays

**DOI:** 10.1101/2025.09.17.676825

**Authors:** Jacob E. Labovitz, Patrick Kulaga, Eric M. DuBois, Kunyu Li, Timothy M. O’Shea

## Abstract

Traumatic injury to the healthy central nervous system (CNS) causes mechanical tissue damage that results in localized cell death and blood-brain-barrier (BBB) disruption. CNS tissue damage stimulates a multicellular wound response to limit the extent of damage but fails to reestablish the normal function of injured tissue. There is strong interest in developing new strategies to augment regeneration after CNS injury. To enable therapy development, reliable assays to screen and identify molecular approaches to augment glial-based wound responses over fibrotic scarring are needed. Scratch assays, which involve mechanically removing cells from an *in vitro* culture, allow for the simulation of wounds with high throughput and tight control over applied treatments to mechanistically study cell migration and proliferation functions that are critical to effective repair. Current methods require researchers to individually scratch each well with a pipette tip, resulting in low throughput as well as inconsistent scratch widths, straightness, and efficacy within and between wells. Here, we describe the design of a quickly assembled (<30min), inexpensive (<$110) scratch assay rig that readily creates uniform scratches that are straight (average tortuosity < 1.1), have tunable widths (730-1100µm), and fully remove damaged cells from the simulated wound region. Designed for a 24-well plate, the rig allows for high-throughput screening of varied experimental conditions or for testing many replicates. Application of the scratch assay device on an *in vitro* culture of neural progenitor cells (NPC) demonstrates the ability to detect differences in wound closure rate for three unique media conditions. These results support the implementation of this high-throughput scratch assay rig as a method to standardize and improve the efficiency of *in vitro* wound healing studies.

## 1. Introduction

Traumatic injury to the central nervous system (CNS), such as traumatic brain injury (TBI) or spinal cord injury (SCI), occurs when neural tissue is damaged by an applied external force or penetrating object. The primary injury is a mechanical insult that results in widespread and indiscriminate cell death, leading to disruption of neural circuits, compromised blood-brain-barrier (BBB), as well as acute hemorrhage and edema [1–4]. In response to injury, the activation of local glial cells and recruitment of peripheral immune and stromal cells functions to clear cellular debris, combat infection, and seal off the wound environment [1,4]. In natural CNS wound repair there is a coordinated interplay between fibrotic and glial cell-based repair mechanisms, with the relative proliferation and migration capacity of the cells involved determining the dominant repair phenotype [5,6]. Typically, in injury to the mammalian adult CNS, high stromal cell and macrophage activity during the wound remodeling phase results in lesions composed of collagen-rich fibrotic scar that affords limited functional recovery [1,5,6]. However, in certain contexts, such as in the mammalian neonate, an elevated proliferation and migratory capacity of astrocytes and microglia drives glia-based repair, resulting in cellular frameworks that support axonal regrowth and neural circuit reconstruction [7]. Glial cell proliferation and migratory capacity is spatially and temporally limited in adult injuries, such that fibrotic scarring is favored at the lesion core and the glial response merely contributes to reinforcing the interface between this forming fibrotic lesion and adjacent preserved neural tissue [5,6]. While these natural CNS wound repair processes have been widely characterized *in vivo* [1,8], the molecular regulation that dictates proliferation and migration competency in glial cells and consequently the magnitude of the glia-based repair relative to fibrotic repair are not well understood. Identifying cell specific approaches to regulate these mechanisms will be critical for developing strategies to enhance endogenous glia-based repair following injury. Therefore, there is a need to implement reproducible, high throughput assays to dissect cellular responses to mechanical damage and test therapeutic strategies to enhance neural regeneration-determining mechanisms.

Scratch assays are commonly used *in vitro* to study wound repair mechanisms of cells from a variety of different organ systems in a tightly controlled environment [9]. Classically, these assays involve culturing a monolayer of cells and scoring the cells with an apparatus such as a pipette tip, damaging cells and creating a linear, acellular region or a ‘wound’ within the monolayer. The primary readout of this assay is wound closure as evaluated by microscopy (**Fig. 1A**), with the rate of closure being a function of both cell proliferation and migration [9,10]. By altering cell media composition or introducing molecular regulators such as protein growth factors or small molecule drugs, this assay enables mechanistic investigation of the key drivers of these two cellular processes that can be used to formulate strategies to regulate and enhance repair outcomes after tissue injury. Scratch assays are accessible, being both lower cost than *in vivo* models and simple, requiring minimal equipment or established assay expertise [9]. Common techniques for creating scratches include manual scoring with pipette tips or cell scrapers, or using molds to prevent cell growth in defined regions [9–11]. While these approaches are accessible and low-cost, the manual scratch methods employed rely on precise training of personnel, which can make them prone to variability in scratch geometry. Variations in scratch width and edge roughness can mask the effect size of therapeutic interventions and hamper mechanistic interpretation of factors that modulate wound closure outcomes. Meanwhile, mold-based approaches yield highly uniform gaps but do not produce mechanical injury or chemical cues to stimulate wound healing [11,12]. These manual assays also have limited throughput as they are often performed in 6- or 12-well plate configurations, limiting experimental replicates and making screening larger libraries of potential therapeutic molecules labor intensive [11,12].

**Figure 1:**
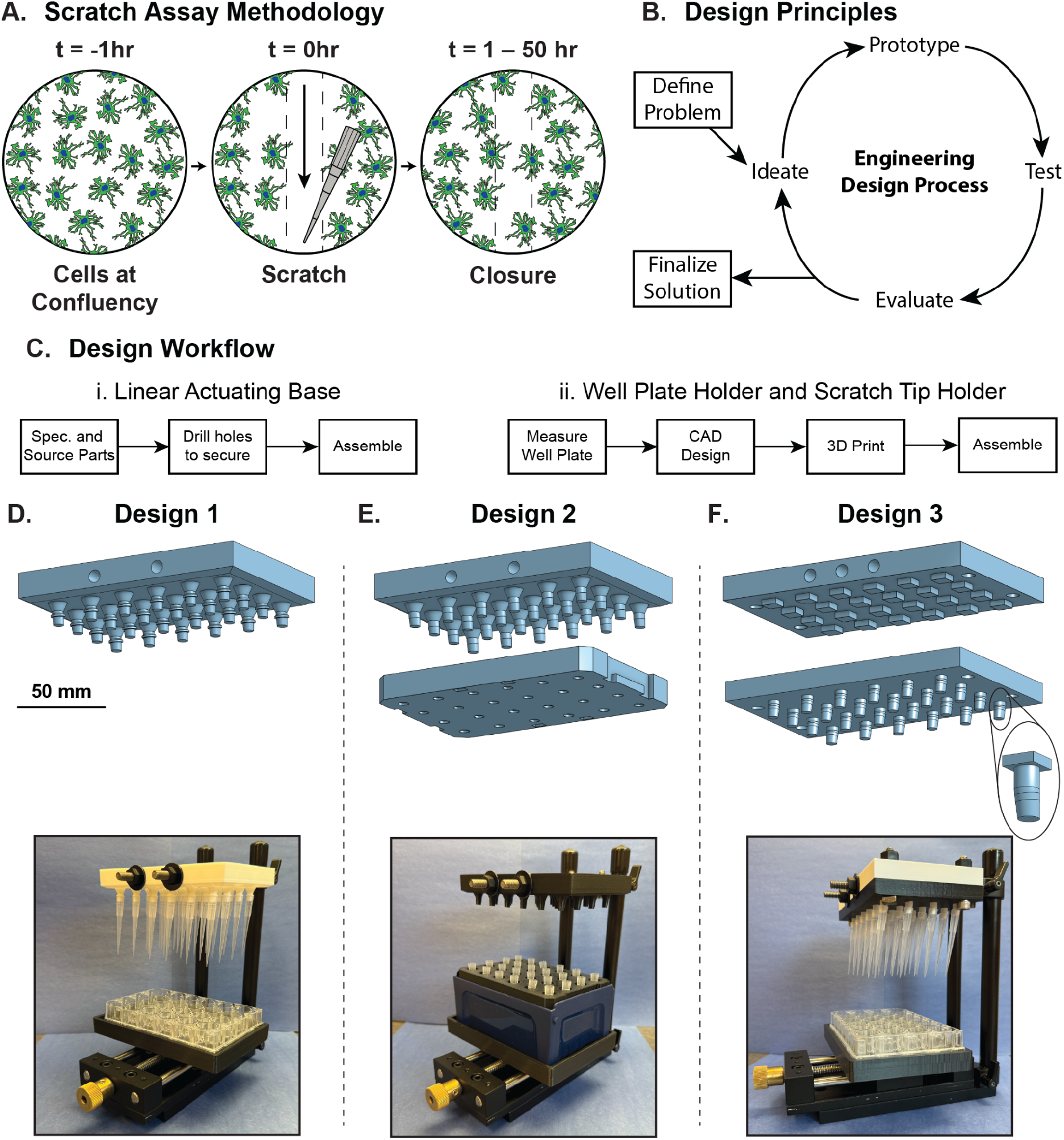
Design Process and Iterative Prototypes. **A**. Schematic demonstrating the traditional scratch assay methodology. **B**. Flow diagram highlighting the core principles of the engineering design process implemented in the design of this device. **C**. Flow diagram illustrating: (i) the steps to create the linear actuating base and (ii) the well plate holder and scratch tip holders. **D**. Above: CAD rendering of Scratch Tip Holder - Design 1. Below: Image of fully assembled Design 1 Scratch Rig. **E**. Above: CAD rendering of Scratch Tip Holder -Design 2 and the pipette tip rack required for loading tips. Below: Image of fully assembled Design 1 Scratch Rig. **F**. Above: CAD rendering of Scratch Tip Holder - Design 3, including the Cap Plate (top), Shaft Plate (bottom), and Tip Holder (detail image). Below: Image of fully assembled Design 1 Scratch Rig.

Several device-based solutions have been proposed to address the limitations of manual scratch methods. For example, some groups have developed patterned polymer stamps (ie. polydimethylsiloxane), which can cause small regions of cell damage through contact-induced mechanical compression, but these designs are still low throughput and may not entirely remove cells from the wound area [13,14]. Laser-based ablation techniques have been applied to precisely create wounds, but are prohibitive as they require specialty equipment to implement [10,15]. Manual multichannelled scratchers have been developed to improve the throughput, with 8- and 96-armed devices used to create wounds across wells on 96- and 384-well plates respectively [16,17]. These tools require specialized fabrication techniques or parts that are not widely available making them difficult to assemble. Additionally, these techniques still require significant handling, making the scratches prone to high interscratch and interuser variability. Other devices have been developed to address these challenges, but suffer from a range of issues, including: plate format restrictions (ie. 12-well plates or petri dishes), lack of open-source availability, absence of modular or modifiable features, or cost and complexity of assembly [18– 20].

Here, we report the design and fabrication of an open-source scratch assay device that has the capacity to be scaled from 6-well to 96-well plates. All components can be either purchased commercially or 3D-printed using entry level, rapid prototyping machines, enabling widespread accessibility and customization. Iteration on three unique designs led to the development of an optimized device that reproducibly generates scratches across all tested wells of the plate and minimal variation between plates, while maintaining consistent width, low tortuosity and minimal roughness at scratch edges. Relative to manual scratches, the optimized device significantly improved assay throughput and decreased the overall variation in tortuosity and roughness. We tested the device using cultured neural progenitor cells that have distinctly different proliferation and migratory capacity after scratch when cultured under different media conditions, demonstrating that the high throughput scratch set up can readily distinguish differences in wound repair capabilities. This device is an important step towards establishing a high throughput and standardized assay for assessing glial cell responses to mechanical damage *in vitro* and will become an essential and widely used tool as part of our work to develop new mechanistic-based approaches to augment glial cell proliferation and migratory capacity to improve repair outcomes after CNS injury.

## 2. Materials & Methods

### Device Materials

All materials for the scratch assay device can be found in **Table 1**. Product numbers, prices, and URLs to items can be found in **Table S1**. Items 9-13 are commonly found in makerspaces and workshops, further reducing costs. Items 15-18 are 3D printed, as denoted by italics. Files for 3D printed parts are in.STL format and are available upon request.

**Table 1.**
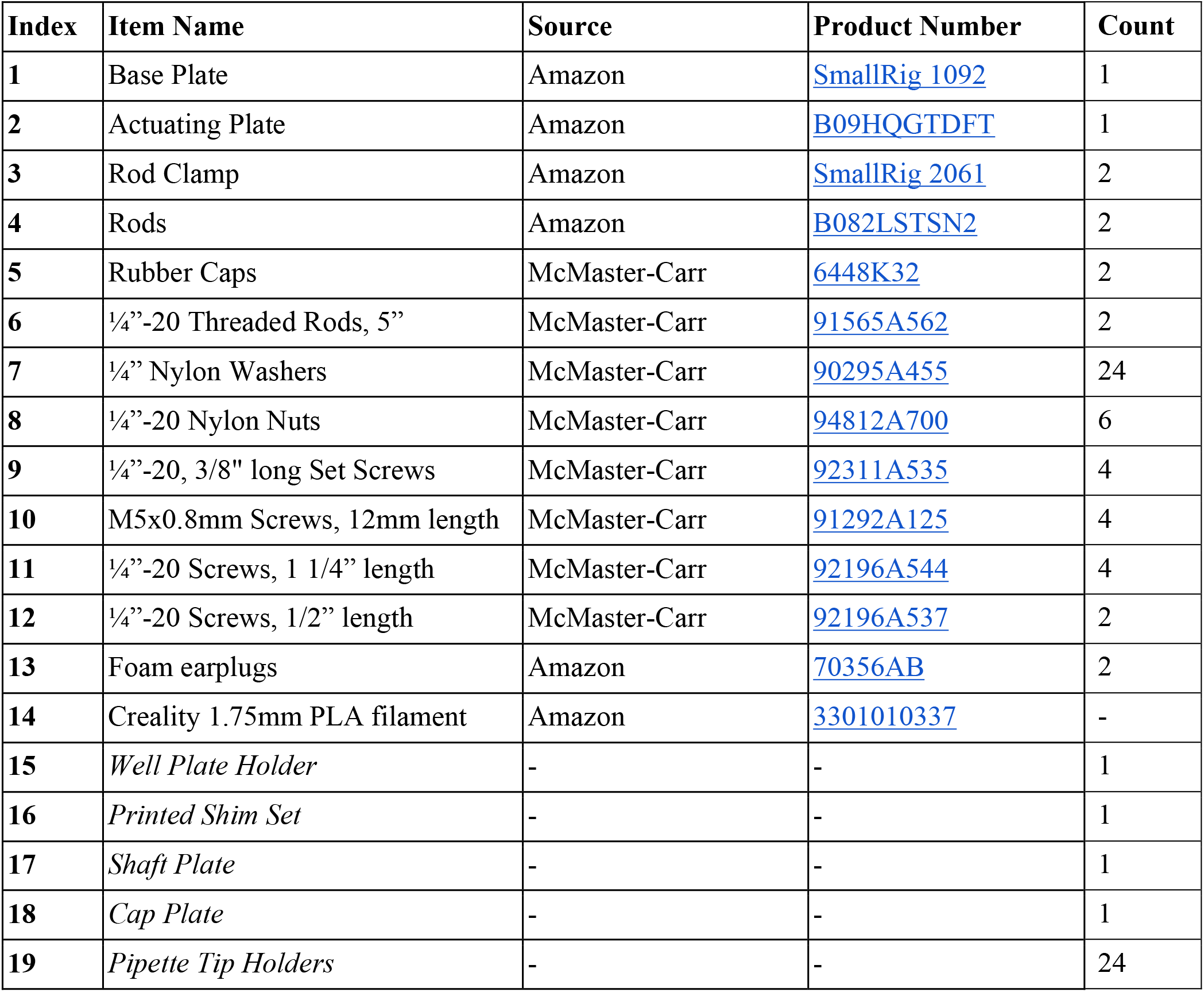
List of all materials for device design

### Computer-Aided Design and 3D Printing

All design modeling was performed using Onshape, a web-based Computer-Aided Design (CAD) platform. CAD drawings for the 24-Well Plate Holder and Design 3 have been provided in **Fig. S1 and Fig. S2**. Models were exported as STL file format and sliced in UltiMaker Cura. Slicing parameters were set to 0.1mm layer height, 5 layer (0.5mm) top and bottom thickness, and 20% gyroid infill.

All models were printed using Creality 1.75mm polylactic acid (PLA) filament (**Item 14, Table 1**). PLA filament was selected due to its low cost, ease of use, and compostability. 3D printing was performed using a Creality Ender 3 fused deposition modeling (FDM) device with a bed temperature of 55ºC and printing speed of 50 mm/s with a 0.4mm diameter nozzle at 200ºC. A textured print bed and reducing top/bottom print speed by 20% were sufficient to counteract warping on larger parts, when needed.

### Scratch Assay Rig Assembly and Calibration

The scratch assay rig can be assembled as follows. These steps have been recorded as a video and are available upon request. Hand tightening is sufficient for any steps using **nylon nuts** (**Item 8, Table 1**).

#### Tools Needed

- ⅛” Allen Wrench
- 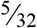” Allen Wrench
- 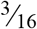” Allen Wrench
- Level
- Square
- Razor blade
- Drill Press
- 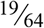” Drill Bit

### Optional

- 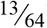” Drill Bit
- ¼”-20 Tap

#### Base Sub-assembly

*Note: all bolded references in the assembly section (i*.*e*. ***(2)****) are references to Items in* ***Table 1***

1. Ream the countersunk holes on the base of the **actuating plate (2)** using a 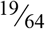 ” drill bit.
2. Orient the base plate such that the four untapped holes align with the knob on the actuating plate. This row of untapped holes will be referred to as the bottom.
3. Join the **base plate (1)** and **actuating plate (2)** with a **½” screw** (**12)**. Use a square to ensure proper alignment.
  a. While the friction from a single screw should be sufficient to prevent motion, a second hole on the bottom right of the base plate can optionally be drilled and tapped with a 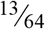 ” drill bit and ¼”-20 tap, and joined to the actuating plate with another ½” screw (12).
4. Join **1 rod clamp (3)** to the **base plate (1)** using the provided ¼”-20 x ¾” screws.
5. Insert **both aluminum rods (4)** into the attached **rod clamp (3)** such that the end of the rods are flush with the base plate. Clamp the rods in place.
6. Slide the remaining **rod clamp (3)** onto the **aluminum rods (4)** and clamp in place. Cap the exposed end of each rod with a **rubber cap (5)**.
7. Attach the **well plate holder (**15) to the **actuating plate (2)** stage using **4 M5×0.8 screws (10)**.
  a. Use a leveler to ensure the well plate holder is level. The **printed shim set (16)** can be used to adjust tilt until level.
8. Screw **2 threaded rods (6)** into the outermost threaded holes on the upper **rod clamp (3)**, such that they are parallel to the base.
9. Add **4 set screws (9)** to lock the **threaded rods (6)** in place. Place 1 set screw above and below each rod.
  a. These set screws can be used to adjust the tilt of the mechanism. Loosen the bottom set screws and tighten the top set screws to tilt the front of the mechanism up. Loosen the top set screws and tighten the bottom set screws to tilt the front of the mechanism down.

#### Mechanism Sub-assembly

10 Place **1 pipette tip holder (19)** into each shaft of the **shaft plate (17)**. Orient the tip holders such that the angle will point towards the knob of the actuating plate.
11 Cut 2 **foam earplugs (13)** into 24 pieces using a razor blade such that each piece fills the remaining space in a shaft.
12 Fit the **cap plate (18)** onto the **shaft plate (17)**. Thread **4 1 ¼” screws (11)** through the aligned holes to join the sub-assembly. Fasten with **4 nylon nuts (8)**.
  a. The screws must be oriented with the screwhead facing down to prevent interference with the tip holder mechanism.

#### Final assembly

13 PSlide **11 nylon washers (7)** onto each **threaded rod (6)**.
  a. These washers act as spacers, and more can be used if preferred.
14 Slide the mechanism sub-assembly onto the **threaded rods (6)**. Fasten with **1 nylon washer (7)** followed by **1 nylon nut (8)** per threaded rod.

### Scratch Assay Device Validation

The device was tested using a dry erase marker to simulate a cell monolayer, allowing for efficient testing using 24-well plates (recorded as a video and available upon request). Dry erase was applied in a uniform layer to each well of the plate. Scratches were performed, the scratch was imaged with brightfield microscopy. Pipette tips were replaced to simulate individual tests and replicates.

### Dry Erase Marker Image Analysis

The image processing of dry erase marker scratches has been recorded as a video and is available along with the corresponding code upon request. Scratch images were segmented, binarized, and skeletonized using ImageJ. Scratch length was obtained using the Feret’s Diameter measurement in ImageJ. Tortuosity, average width, and average roughness were computed using MATLAB scripts. Tortuosity was measured as the ratio of the path length of the scratch centerline to the linear distance between its endpoints. Average roughness was calculated as the mean absolute deviation from the line of best fit at both scratch edges.

### Cell Culture and *in vitro* Scratch Assay

Neural progenitor cells (NPCs) used were derived from the neural induction and expansion of mouse embryonic stem cells using procedures outlined in detail previously [21]. NPCs were cultured as previously described [21] and all cell stocks used were maintained below P30. Briefly, NPCs were grown in Neural Expansion (NE) media consisting of Advanced DMEM/F12 with 2% V/V B27 (no Vitamin A) (50X) (Cat# 12587010, ThermoFisher Scientific) and growth factors EGF (Cat# AF-100-15-100UG, Peprotech) and FGF (Cat# 100-18B-100UG, Peprotech) (100 ng/mL each). After 1-2 days of growth, NPCs were processed for plating. To plate, cells were trypsinized, collected, and counted with a hemocytometer after Trypan blue staining. Cells were then resuspended in NE media, gently pipetted for 5-10 minutes to ensure complete dispersion, and transferred to a 24-well plate (Corning Costar, Ref# 3524) at a concentration of ∼1 × 10^5^ cells/well.

NPCs were allowed to adhere for 24hrs in NE media conditioned with either 100 ng/mL E/F (EGF/FGF), 2% fetal bovine serum (FBS), or 50 ng/mL E/F + 2% FBS. To ensure sterility, the scratch assay rig assembly was cleaned with 70% ethanol before being placed in a sterile, laminar-flow bench and exposed to germicidal UV-C for 15 minutes. After sterilization, 200µL pipette tips were pressed onto the pipette tip holders and the cells were scratched using the scratch rig. Subsequently, the wells were rinsed with 1x phosphate buffered saline (PBS) and the conditioned NE media was replaced. The wells were then imaged on an inverted brightfield microscope (Olympus IX83) at ∼12hr intervals up to 48hrs, with a final timepoint at 54hrs. Scratches were analyzed using ImageJ and custom MATLAB code.

### Images Analysis of *in vitro* Scratches

The image processing of *in vitro* scratches has been recorded as a video and is available along with the corresponding code upon request. Images were cropped into squares and processed using a top-hat filter with a radius of 15 pixels, followed by Li’s threshold in ImageJ. Objects less than 1000 pixels in area were then removed. Binarized images were then imported into MATLAB for scratch detection and measurement. Scratches were detected by splitting images in half down a column of pixels devoid of cells. If no such column existed, the midline of the image was used. The edge of each scratch was found by tracing a curve along cells closest to the midline at each height of the image. The resulting curves were then smoothed using a two-point moving average until the second derivative was less than or equal to 0.5.

## 3. Results

### 3.1 The Engineering Design Process yields iterative designs that optimize the number of scratches, ease of use, and variability

We implemented the design engineering workflow to develop a new device capable of producing uniform, consistent scratches while remaining simple to operate (**Fig. 1B**). This process incorporated six steps: (**1**) Defining the problem/scope, (**2**) Ideating solutions, (**3**) Prototyping solutions, (**4**) Testing solutions, (**5**) Evaluating performance, and (**6**) Iterating back to (2) or Finalizing the solution. We defined the scope of the scratch assay requirements using quantifiable design criteria (**Table 2**), which served as clear performance targets to enable decision-making throughout the design process.

**Table 2.** Design criteria of scratch assay device

| Design Criteria: | Justification |
| --- | --- |
| **1)** Scratches must be a minimum of 2mm long and 500µm wide across all wells | Enables reproducible, high throughput results. Window length is larger than 10x microscope field of view (FOV) and width is >10 cells |
| **2)** Intra- and inter-plate variability in width and roughness must be reduced by 40% relative to manual scratches | Enhances reproducibility of results relative to the manual technique |
| **3)** Inter-user variability in width and roughness must be less than 15% the total measurement | Enhances reproducibility of results relative to other users, reducing reliance on a single user for consistent data collection |
| **4)** The device must decrease the time to scratch by >75% | Increases assay throughput, enabling many wells to be tested simultaneously and favors user adoption |
| **5)** The device must cost less than \$200 and be able to be assembled/disassembled in <30 min | Ensures that the device is accessible and can be implemented by any lab, which favors user adoption |
| **6)** The design must be chemically and mechanically resilient and easily repaired | Enables long-term, repeatable use, which favors user adoption and accessibility |
| **7)** Does not need sterilization after each use | Limits prep time before scratch, which favors user adoption |

To achieve these design criteria, we entered the second step of the design engineering workflow: (**2**) **Ideating** solutions. Our initial concept used a hybrid approach in which some components were commercially purchased and others were 3D printed in-house. A low-cost, linear actuating base was selected to avoid designing all parts *de novo* and to eliminate potentially costly machining. We next opted to 3D print a 24-well plate holder, which was designed to secure a tissue culture (TC) plate on the linear actuating base (**Fig. S1**). While 3D printing resources are not universally available, they are increasingly accessible at academic institutions, makerspaces, and through low-cost commercial services (such as Protolabs Network or Makelab). Together the actuating base and 24-well plate holder formed the core of the scratch assay rig (**Fig. 1C**), providing a stable foundation for the iterative development of the scratch tip holder.

When ideating the design of the scratch tip holder, we selected 3D printed designs to enable rapid, low-cost fabrication and customization. This also ensures low cost replacement of parts if they become contaminated or break. The rig must apply mechanical injury by disrupting cell membranes, which ultimately restricted the design to scratch-based methods that require a cell-device interface. Therefore, we chose to design a device that used standard pipette tips because micropipettes are ubiquitous in academic laboratories, their disposable tips eliminate the need for device sterilization, and the tips are inexpensive and easily replaced compared with 3D-printed or machined interfaces.

Having determined these key features, we (**3**) **Prototyped** the proposed solution. Using Onshape software, we designed a scratch tip holder that could fit 200µL pipette tips via a press fit connection at a fixed angle. To (**4**) **Test** this solution, we used a dry erase marker-based scratch assay (**Fig. S3**), enabling high throughput testing with minimal expense. On (**5**) **Evaluation**, the first design failed our criteria as it could not scratch all wells simultaneously and the fixed 90° angle tip caused skipping, producing discontinuous scratches. To overcome these identified concerns, we (**6**) **Iterated** this design, adding leveling shims and set screws into the plate design, angling the pipette tip holders to 98° (+8° from vertical), and adding ridges to secure the pipette tips. These design changes culminated in the Design 1 scratch tip holder (**Fig. 1D, Table 3**).

**Table 3.** Features of each Scratch Assay Rig design iteration.

| Design 1 | Design 2 | Design 3 |
| --- | --- | --- |
| 1. Scratch plate requires single print<br>2. 24-well plate design<br>3. Angled pipette tip holder<br>4. Ridges hold pipette tips in position | 1. Scratch plate requires two prints<br>2. 24-well plate design<br>3. Angled pipette tip holder<br>4. No ridges to hold pipette tips in position<br>5. Pipette Tip Rack to set the height of tips | 1. Scratch plate requires multiple prints<br>2. 24-well plate design<br>3. Angled pipette tip holder<br>4. Readily customizable components<br>5. Modular<br>6. Foam spring between plates for movement in z-direction |

While Design 1 was able to scratch many of the wells in a high-throughput manner, it suffered inter user variability in success rate, with different lab members achieving varying levels of success. Additionally, we observed that there were differences in scratch success and scratch morphology on an inter-plate basis. We hypothesized that this was a function of the variation in pipette to well plate contact due to the high geometric tolerances inherent to the 3D print, resulting in unequal contact/applied pressure. Stereolithography printing was briefly tested due to its tighter tolerances, however cost and accessibility concerns led to our continued focus on FDM printing. After evaluating this outcome, we iterated a new prototype.

Design 2 (**Fig. 1E, Table 3**) removed the tip-leveling ridge, instead using a pipette tip rack to mount all pipette tips evenly onto the scratch device. We hypothesized that this would help overcome variations in tip length and 3D print layer height that result in missed or disrupted scratches. Testing this prototype, we observed an increased rate of missed wells between users, greater variability in scratch geometry, and found that users took longer to scratch as a result of the tip holder. Additionally, in both Design 1 and Design 2, we faced significant 3D printing challenges, including warping of the flat scratch tip holder and filament deposition errors on the angled tips.

To overcome these challenges, we created a third iteration of the scratch tip holder, which featured a modular design (Design 3). This design allowed for the tips to be printed separately from the flat scratch tip holder, decoupling the concerns of part warping from filament deposition errors. An added benefit of this design is that the tip holders could be mass produced, allowing for substitution if they are damaged and for the design to be rapidly altered if required. This Design 3 scratch tip holder (**Fig. 1F, Table 3**), also featured a plate that allowed for the individual tip holders to translate in the z-direction via foam inserts. This design change allowed for all tips to contact the well plate under applied pressure, overcoming variability across users and scratch morphology. Having iterated three unique scratch plate designs, we then sought to evaluate these designs using the quantitative design criteria.

### 3.2 Design 3 Scratch Rig creates scratches with consistent width and roughness

We next evaluated the performance of the 3 designs for the critical quantitative metrics outlined in the design criteria (**Table 2**), including: scratch width, length, roughness, and variability. To permit high-throughput and low cost testing of Scratch Rig designs, we tested scratch rig performance on 24-well plates coated with dry erase marker, which, when allowed to dry, forms a thin film composed of black pigment, resin binder, and a lubricating silicone-based polymer (**Fig. 2A, S3**). All 24 coated wells within a single plate were scratched using the 3 designs, imaged by a brightfield microscope, and then scratch metrics were quantified using custom-made image analysis code. Scratching was repeated across three separate plates for each rig design. We evaluated the scratch success rate, length, and tortuosity (path length/linear distance between endpoints) across the entire scratch but assessed scratch width and roughness within a restricted field of view (FOV) within the center of the scratch in a manner consistent with standard scratch assay quantification approaches used in the field (**Fig. S3**). Scratch success rate for a single user across three 24-well plates was as follows: Design 1 (71/72 wells), Design 2 (70/72 wells), and Design 3 (72/72 wells), demonstrating a slight improvement in reproducibility for Design 3 (**Fig. 2B**). Average scratch lengths ranged from 10.2 mm (Design 3) to 11.1 mm (Design 1) (**Fig. S4**), which are all sufficiently large to select a 2 mm FOV for width and roughness quantifications. Average plate tortuosity ranged from 1.06 to 1.08 for all designs, indicating that all designs create scratches with less than 10% deviation from linearity (tortuosity of 1), making it practically indistinguishable from a straight line (**Fig. S4**).

**Figure 2:**
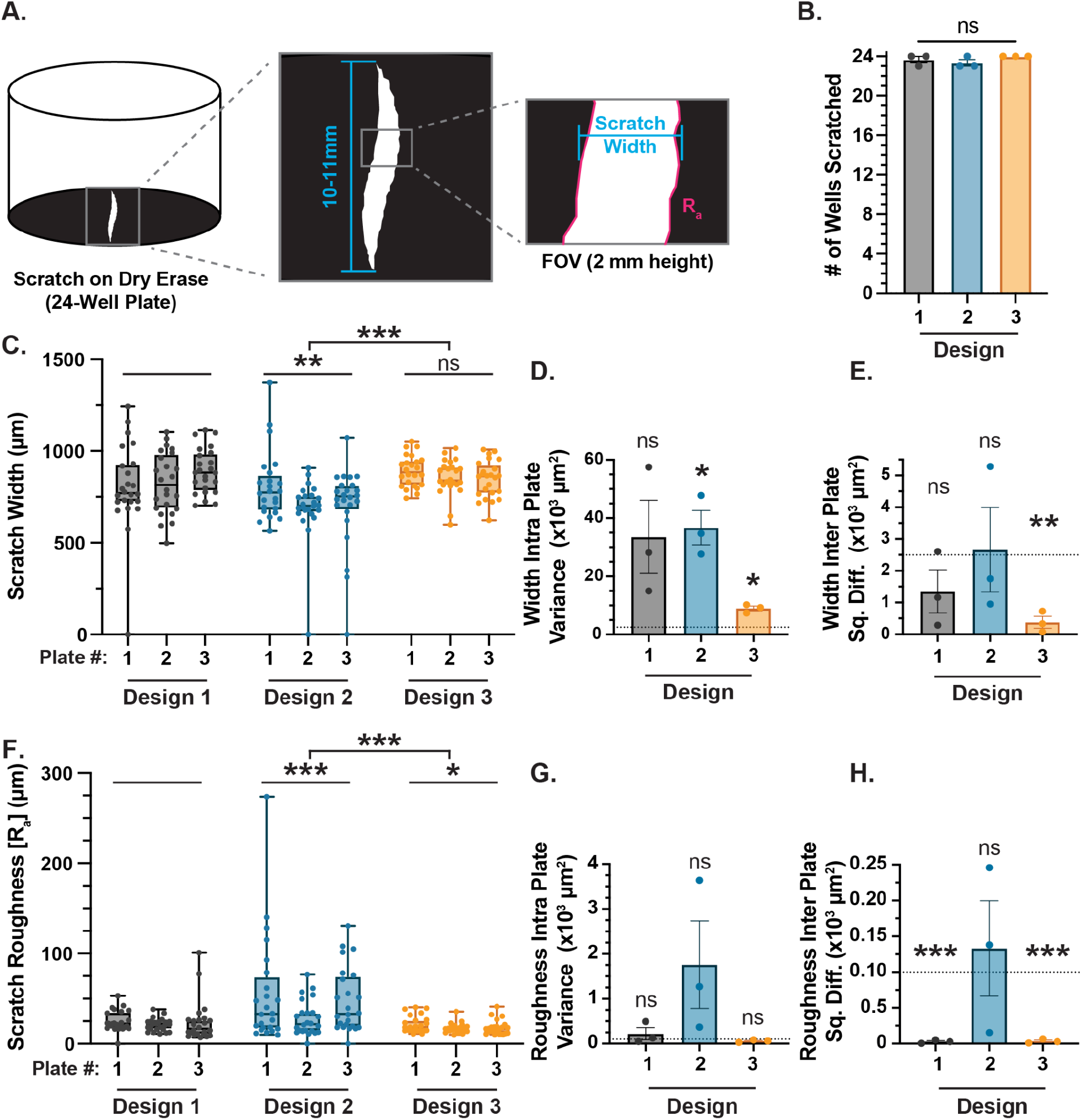
Iterative design of scratch assay rig optimizes number of scratches, ease of use, and limits variability. **A**. Diagram of dry erase scratch assay highlighting scratch width and average roughness (R_a_) measurements. **B**. Bar graph showing the effect of design iteration on the number of wells scratched on a 24-well plate. Not significant (ns) across all conditions, one-way ANOVA with Tukey’s multiple comparison test. Graph shows mean ± s.e.m. with individual data points showing n = 3 plates. **C**. Box and whisker plot showing the effect of design on scratch width (µm). Box shows mean and interquartile range (IQR), whiskers show full data range with individual data points showing n = 24 wells per plate. **D**. Bar graph showing the effect of design iteration on the intra-plate variance for scratch width. Graph shows mean ± s.e.m. with individual data points showing n = 3 plates. **E**. Bar graph showing the effect of design iteration on the inter-plate squared differences (Sq. Diff.) for scratch width. Graph shows mean ± s.e.m. with individual data points showing n = 3 plates. **F**. Box and whisker plot showing the effect of design on scratch roughness (µm). Box shows mean and interquartile range (IQR), whiskers show full data range with individual data points showing n = 24 wells per plate. **G**. Bar graph showing the effect of design iteration on the intra-plate variance for scratch roughness. Graph shows mean ± s.e.m. with individual data points showing n = 3 plates. **H**. Bar graph showing the effect of design iteration on the inter-plate squared differences (Sq. Diff.) for scratch roughness. Graph shows mean ± s.e.m. with individual data points showing n = 3 plates.

Design 3 produced consistent scratch width values with an average of 864 µm across 3 plates (24 wells each) (**Fig. 2C**). Design 1 was more variable, but generated a similar average scratch width of 830 µm (**Fig. 2C**). Design 2 showed the greatest variability and a significantly smaller average scratch width of 733 µm (**Fig. 2C**). Variance in the scratch width parameter across the 3 designs was quantified within plates (i.e. between the 24 wells on a given plate) to derive intra-plate variability (σ^2^, **Eq. 1; Fig. 2D**) and between plates (i.e. relative to the population mean) to derive inter-plate distance from mean (**Eq. 2; Fig. 2E**). To assess successful mitigation of variability we created threshold values of acceptable variance values of 2,500 µm^2^ for scratch width (50µm standard deviation) and 100 µm^2^ for scratch roughness (10 µm deviation). Design 3 achieved scratches with the least intra-plate variability, relative to Designs 1 and 2, but this value was still significantly greater than our proscribed threshold. Design 3 showed the least inter-plate variability relative to Designs 1 and 2, and this variability was within our designated threshold, suggesting that the design achieved reproducible scratches across plates but that there was still some significant variability across individual wells within a given plate.

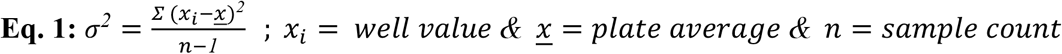

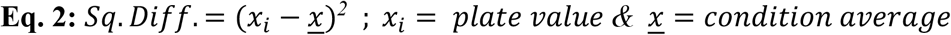

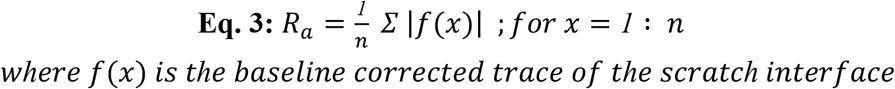

Design 3 also achieved scratches whose average roughness (R_a_, **Eq. 3**) was the smallest across the rig designs, with an R_a_ of 18.8 µm across the FOV, equating to approximately 1-2 cells wide (**Fig. 2F**). Design 1 had a similar average roughness of 23.7 µm, but Design 2 created significantly rougher scratches, with an average value of 45 µm (**Fig. 2F**). Assessing the intra-(**Fig. 2G**) and inter-(**Fig. 2H**) plate variability revealed that Design 1 and Design 3 perform similarly, with less overall variation than Design 2. All 3 designs demonstrate no significant difference between the mean value and the roughness threshold of 100 µm^2^ for intra-plate variability, but Design 1 and Design 3 were markedly lower than the threshold value for inter-plate mean square difference. These data show that the Design 3 rig can consistently create scratches across all wells with substantially smaller intra- and inter-plate variability in scratch width and roughness, making it a more robust scratch apparatus.

Statistics: For **C** and **F** a Brown-Forsythe and Welch ANOVA test (Gaussian distribution with unequal SD’s) was performed followed by a Dunnett’s T3 multiple comparisons test.

Statistical comparisons to Design 1 or as designated. Not significant (ns), *P < 0.05, **P < 0.005, and ***P ≤ 0.0005 across all conditions. For **D, E, G**, and **H** a one-sample t-test was performed to compare the mean value against a threshold of 2500 µm^2^ (**D, E)** or 100 µm^2^ (**G, H**). Not significant (ns), *P < 0.03, **P < 0.008, and ***P ≤ 0.0004 across all conditions.

### 3.3 Optimized rig is highly reproducible and can be tuned

Comparing all three designs, we selected Design 3 for further characterization and use due to its 100% scratch success rate, low tortuosity, and minimized variability for the scratch width and roughness parameters. To assess the reproducibility and tunability of the Design 3 Scratch Rig, we tested four key performance variables: the user (**Fig. 3A**), the scratch speed (**Fig. 3E**), the outer diameter of the scratching pipette (**Fig. 3I**), and the pipette angle during scratching (**Fig. 3M**). Since changes in scratch length and tortuosity varied minimally, we focused on scratch width and roughness assessments for the four performance variables.

**Figure 3:**
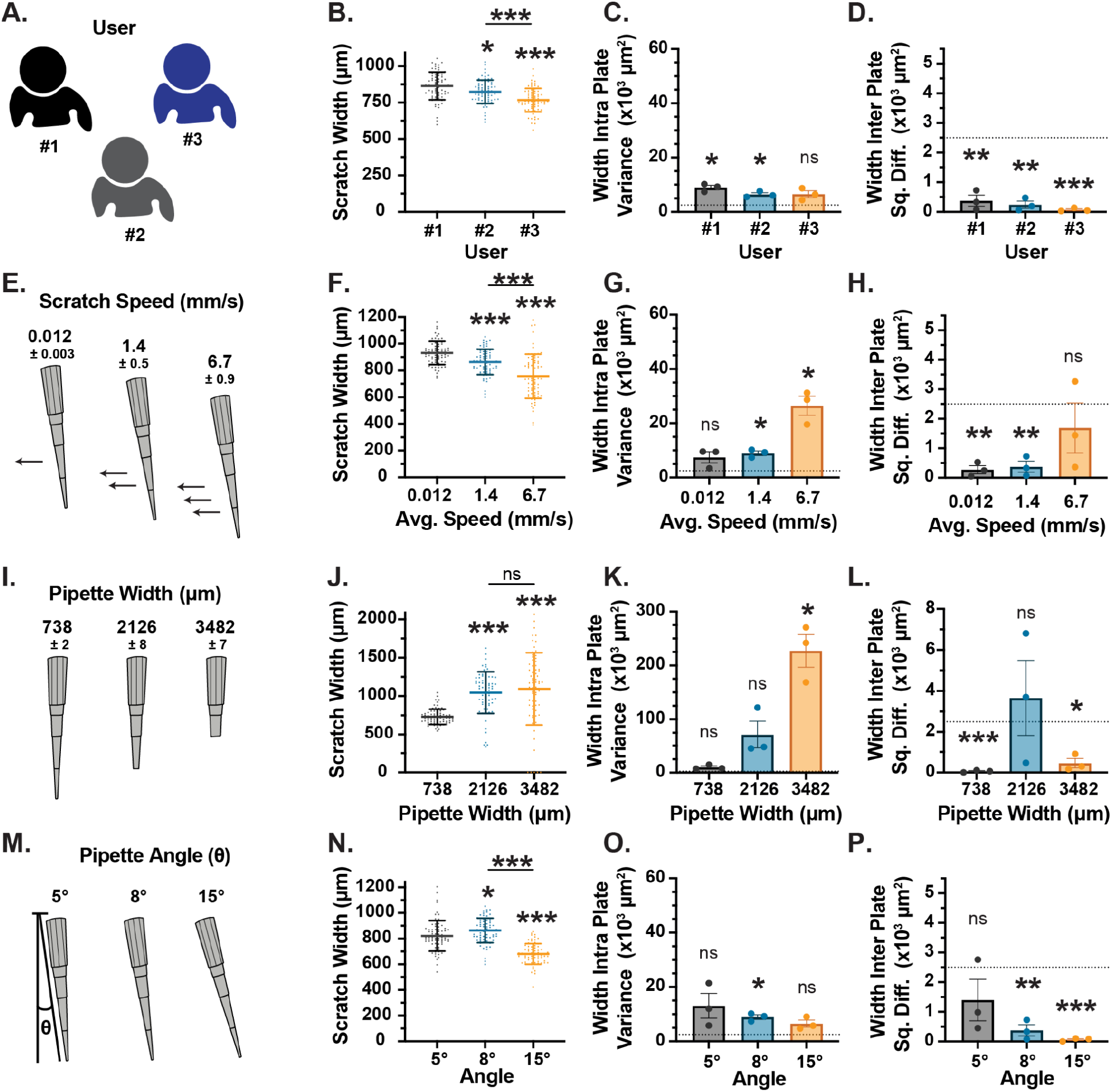
Design 3 scratch rig creates controlled scratches that can be tuned using simple tip parameters. **A**. Schematic showing tested variable: User. **B**. Graph showing the effect of user on scratch width. Statistical comparisons to User #1 or as designated. Graph shows mean ± SD with individual data points showing n = 72 wells from three 24-well plates. **C**. Bar graph showing the effect of user on the intra-plate variance for scratch width. Graph shows mean ± s.e.m. with individual data points showing n = 3 plates. **D**. Bar graph showing the effect of user on the inter-plate squared differences (Sq. Diff.) for scratch width. Graph shows mean ± s.e.m. with individual data points showing n = 3 plates. **E**. Schematic showing tested variable: Scratch Speed. Reported values are the mean ± the 95% confidence interval (CI). **F**. Graph showing the effect of scratch speed on scratch width. Statistical comparisons to 0.012 (mm/s) or as designated. Graph shows mean ± SD with individual data points showing n = 72 wells from three 24-well plates. **G**. Bar graph showing the effect of scratch speed on the intra-plate variance for scratch width. Graph shows mean ± s.e.m. with individual data points showing n = 3 plates. **H**. Bar graph showing the effect of scratch speed on the inter-plate squared differences (Sq. Diff.) for scratch width. Graph shows mean ± s.e.m. with individual data points showing n = 3 plates. **I**. Schematic showing tested variable: Pipette Width. Reported values are the mean ± the 95% confidence interval (CI). **J**. Graph showing the effect of pipette width on scratch width. Statistical comparisons to narrowest width or as designated. Graph shows mean ± SD with individual data points showing n = 72 wells from three 24-well plates. **K**. Bar graph showing the effect of pipette width on the intra-plate variance for scratch width. Graph shows mean ± s.e.m. with individual data points showing n = 3 plates. **L**. Bar graph showing the effect of pipette width on the inter-plate squared differences (Sq. Diff.) for scratch width. Graph shows mean ± s.e.m. with individual data points showing n = 3 plates. **M**. Schematic showing tested variable: Pipette Angle. **N**. Graph showing the effect of pipette angle on scratch width. Statistical comparisons to 5° or as designated. Graph shows mean ± SD with individual data points showing n = 72 wells from three 24-well plates. **O**. Bar graph showing the effect of pipette angle on the intra-plate variance for scratch width. Graph shows mean ± s.e.m. with individual data points showing n = 3 plates. **P**. Bar graph showing the effect of pipette angle on the inter-plate squared differences (Sq. Diff.) for scratch width. Graph shows mean ± s.e.m. with individual data points showing n = 3 plates.

Beginning with User (**Fig. 3A**), we assessed the scratch outcomes for 3 independent users of the Design 3 scratch rig. Each user was given a single plate to practice on before beginning the test. Scratch widths varied between 767 µm (User 3) and 864 µm (User 1) and were significantly different between users (**Fig. 3B**). Intra-(**Fig. 3C**) and inter-(**Fig. 3D**) plate variability was comparable between all users, with 2 out of 3 users exceeding the intra-plate variability threshold and all three users having significantly lower inter-plate variation relative to the 2,500 µm^2^ threshold. Scratch roughness varied between 16 µm (User 2) to 19 µm (User 1) with no significant differences between users and no significant differences in variability (**Fig. S5**). Intra-plate variability was not significantly different from the threshold value, while all 3 users had significantly lower inter-plate variation relative to the 100 µm^2^ threshold. These data suggest that there may be some limited but detectable inter-user variability, but that within a single user’s set of experiments there is excellent consistency.

We next assessed the effect of scratch speed on the resulting scratch morphology (**Fig. 3E**). Using a slow, normal, and fast linear actuator turning speed, we were able to generate scratch speeds of 0.012±0.003, 1.4 ±0.5, and 6.7±0.9 mm/s (mean±95% Confidence Interval (CI)).

Increasing scratch speed resulted in decreasing scratch width, with slow, normal, and fast speeds yielding scratches with widths 931µm, 864µm, and 757 µm, respectively (**Fig. 3F**). Intra-(**Fig. 3G**) and inter-(**Fig. 3H**) plate variability was greatly increased at high scratch speeds, with intra-plate variability exceeding the designated threshold at normal and fast speeds. Meanwhile, the inter-plate squared difference for the slow and normal speeds was significantly lower than the 2,500 µm^2^ threshold. Roughness also demonstrated significant scratch speed dependency.

Scratching at slow and fast speeds resulted in increased scratch roughness (increases of 5 and 11µm respectively) relative to the normal speed (**Fig. S5**). Similar to scratch width, we also observed a significant increase over the 100 µm^2^ threshold in intra-plate variability at the highest speed, but all three speeds yielded inter-plate variation below the proscribed threshold (**Fig. S5**). These data show that scratch width is inversely related to scratch speed, however fast scratch speeds result in rougher and highly variable scratches.

Pipette outer diameter width was the next variable evaluated (**Fig. 3I**). We achieved different sizes by clipping the pipette tips at predefined locations to achieve 3 distinct diameter values of 738±2, 2126±8, and 3482±7 µm (mean±95% Confidence Interval (CI)) (**Fig. S6**). Increasing the diameter allowed for us to vary the scratch width between 730-1100 µm, with the two wider orifice diameters of 2126 and 3482 µm giving significantly wider scratches relative to the unmodified pipette tip (**Fig. 3J**). With each increase in tip diameter, we observed an increase in intra-(**Fig. 3K**) plate variability with the widest tip orifice giving us variability that exceeds the predefined threshold. Meanwhile, the unclipped and widest pipette orifices generated significantly lower inter-plate variation relative to the threshold (**Fig. 3L**). Roughness significantly decreased for only the widest pipette orifice, with an average roughness of 10.8 µm (**Fig. S5**).There is a significant decrease below the 100 µm^2^ threshold in intra-plate variability for the widest pipette tip, but all three orifice sizes yielded inter-plate variation below the threshold (**Fig. S5**). These data show that increasing pipette orifice diameter can significantly increase scratch width without altering roughness, but exacerbates intra-plate variability.

Finally, we tested the effect of the pipette angle on scratch width and roughness (**Fig. 3M**). From our initial testing, we noted that scratches performed normal to the surface resulted in skipping of the tip across the surface forming a stuttering scratch. Meanwhile, scratches performed at oblique angles to the surface resulted in tip buckling, deformation and ultimately failure to scratch.

Therefore, the designs outlined above all had the pipette tip angled at an 8° deviation from the normal (θ = 8°, **Fig. 3M**), resulting in acute angles between the pipette tip and the well plate surface. To test the effect of small changes in the acute angle, pipette tip holders were printed with pipette tip inserts angled at 5° and 15° deviations from the normal. There was a small but significant decrease in scratch width with the 5° angle (822 µm) relative to the standard 8° (864 µm) while the more acute 15° angle also decreased the scratch width significantly to 681 µm (**Fig. 3N**). Intra-(**Fig. 3O**) and inter-(**Fig. 3P**) plate variability decreased with increasing angle, with only 8° angle having intra-plate variability that was significantly greater than threshold.

Average roughness varied between 17-19 µm, but was not different across all conditions (**Fig. S5**). Intra-plate variability was not altered by tip angle, while inter-plate variability was significantly less than the threshold in all cases (**Fig. S5**). These data show that changing pipette scratch angle can modulate scratch width and decrease inter-plate variability while having minimal effect on roughness.

Collectively, these data show that the scratch assay device creates scratches with reproducible width and roughness under set parameters (tip angle, scratch speed, etc), allowing for the high-throughput production of low variation scratches by any user. By tuning these parameters, one can achieve differences in scratch width and roughness, with adjustment of pipette tip angle allowing for narrow, low variability scratches. By contrast varying the pipette tip outer diameter size is the most effective approach for creating wider scratches but introduces greater variability across wells.

Statistics: For **B** and **N** a one-way ANOVA (Gaussian distribution with equal SD’s) was performed followed by a Tukey’s multiple comparison test. Not significant (ns), *P < 0.04, and ***P < 0.0004 across all conditions. For **F** and **J** a Brown-Forsythe and Welch ANOVA test (Gaussian distribution with unequal SD’s) was performed followed by a Dunnett’s T3 multiple comparisons test. Not significant (ns) and ***P< 0.0001 across all conditions. For **C, D, G, H, K, L, O** and **P** a one-sample t-test was performed to compare the mean value against a threshold of 2500 µm^2^. Not significant (ns), *P < 0.03, **P < 0.008, and ***P < 0.0003 across all conditions.

### 3.4 Optimized rig demonstrates improvement on manual scratch

Having characterized Design 3, we benchmarked the performance of the rig under the standard configuration of 8° angle, standard pipette tips and ‘normal’ speed against the manual scratch technique, which is currently the gold standard approach used for scratch assays. Scratches made by the Design 3 rig were on average 864µm wide, as compared to the significantly smaller 591µm manual scratches (**Fig. 4A**). Intra-plate variability (**Fig. 4B**) decreased with Design 3 compared to the manual method but both were greater than the pre-defined threshold variance, while inter-plate variability (**Fig. 4C**) was maintained below the threshold in both cases. There was no difference in roughness between the two methods, with Design 3 generating scratches with an average roughness of 18.8 µm and manual scratches yielding a roughness of 18.3 µm (**Fig. 4D**). Intra-(**Fig. 4E**) and inter-(**Fig. 4F**) plate variabilities were comparable between manual and rig-based scratches, with Design 3 only improving on inter-plate variability relative to the threshold. With an average tortuosity of 1.25 the manual scratches were significantly less straight than those made with the Design 3 rig (1.07) (**Fig. S7**). Scratches made with the Design 3 rig were minimally variable in tortuosity both within (**Fig. S7**) and between (**Fig. S7**) plates. The scratches made by Design 3 were 10.2 mm in length, which is slightly shorter than the 11.3 mm manual scratches (**Fig. 4G**). The greatest improvement afforded by the Design 3 Rig over the manual scratch was to the time required to complete the scratching of all wells within a 24 well plate set up. Beginning with the tips loaded on the rig, the scratch device took 5.7 seconds to load a plate and complete 24 individual scratches. Meanwhile, manual scratching requires 20x the time, requiring 114 seconds to scratch all 24 wells successfully (**Fig. 4H**). This improvement enhances the throughput of the assay and enables rapid, sterile scratches of 24-well plates.

**Figure 4:**
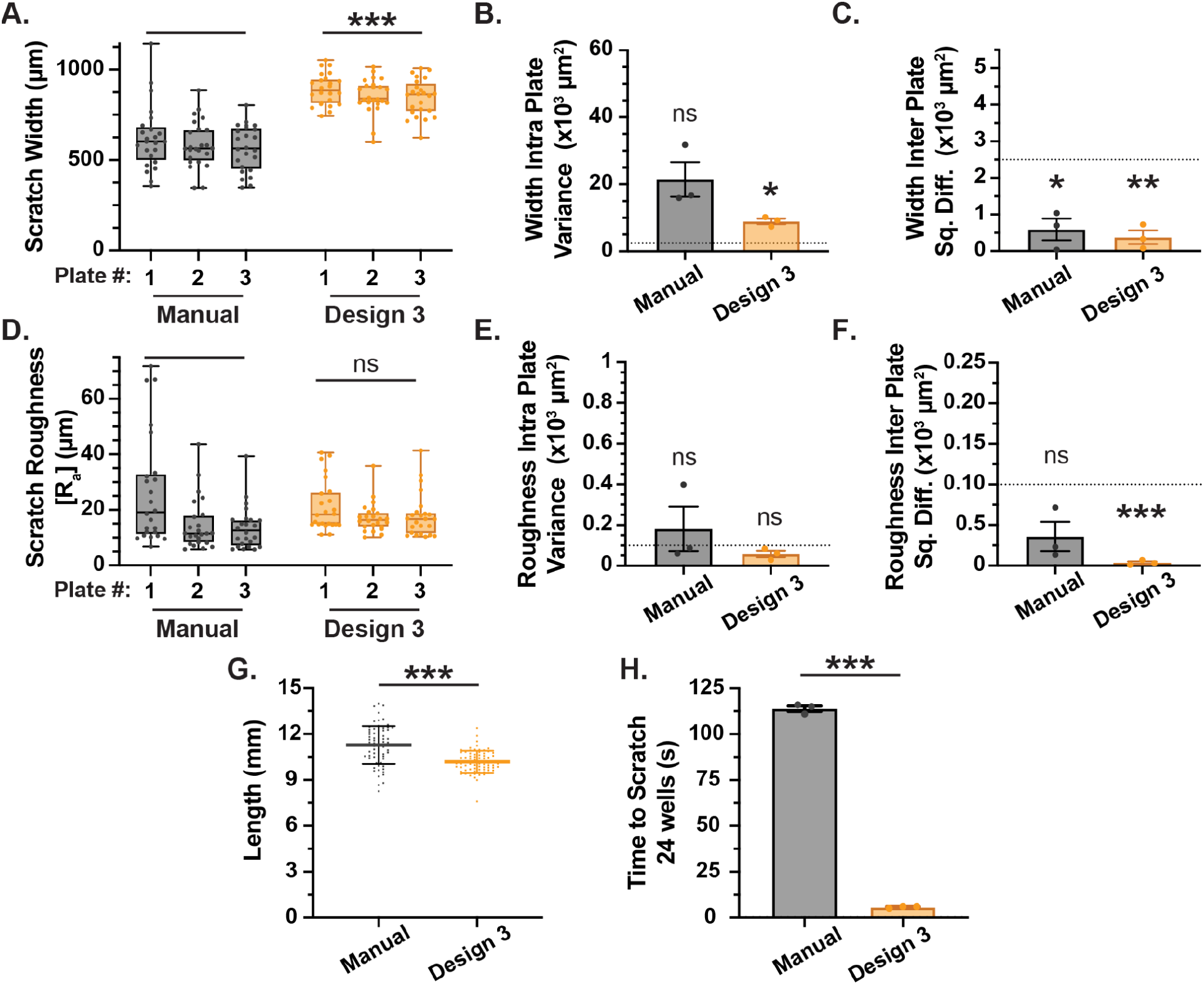
Design 3 Scratch Rig enhances efficiency and consistency when compared with a manual scratch. **A**. Box and whisker plot showing the effect of scratch rig implementation on scratch width (µm). Box shows mean and interquartile range (IQR), whiskers show full data range with individual data points showing n = 24 wells per plate. **B**. Bar graph showing the effect of scratch rig implementation on the intra-plate variance for scratch width. Graph shows mean ± s.e.m. with individual data points showing n = 3 plates. **C**. Bar graph showing the effect of scratch rig implementation on the inter-plate squared differences (Sq. Diff.) for scratch width. Graph shows mean ± s.e.m. with individual data points showing n = 3 plates. **D**. Box and whisker plot showing the effect of scratch rig implementation on scratch roughness (µm). Box shows mean and interquartile range (IQR), whiskers show full data range with individual data points showing n = 24 wells per plate. **E**. Bar graph showing the effect of scratch rig implementation on the intra-plate variance for scratch roughness. Graph shows mean ± s.e.m. with individual data points showing n = 3 plates. **F**. Bar graph showing the effect of scratch rig implementation on the inter-plate squared differences (Sq. Diff.) for scratch roughness. Graph shows mean ± s.e.m. with individual data points showing n = 3 plates. **G**. Graph showing the effect of technique on scratch length. Graph shows mean ± SD with individual data points showing n = 72 wells from three 24-well plates. **H**. Bar graph showing the effect of scratch rig implementation on the time to scratch 24 wells. Graph shows mean ± s.e.m. with individual data points showing n = 3 plates.

In summary, the data presented above demonstrates that we have successfully created a device that fulfills nearly all of our strict design criteria (**Table 2, Table 4**). The Design 3 scratch rig consistently creates scratches that are greater than 2mm long and 500µm wide, fulfilling the first criterion (**Table 4**). Design 3 exceeds the 40% decrease in intra-plate variability in scratch width and inter-plate variability in roughness, but fails to create similar improvements in reproducibility for the reciprocal measurements (**Table 4**). The design rig also limits the variability in the mean scratch width between users to 11% of the total width, but fails to meet the same metric in roughness (**Table 4**). Finally, the device fulfills design criteria 4-7 as it vastly decreases the time to scratch a 24-well plate, is low cost (**Table S1**) and rapidly built, and requires minimal maintenance (**Table 4**).

**Table 4.**
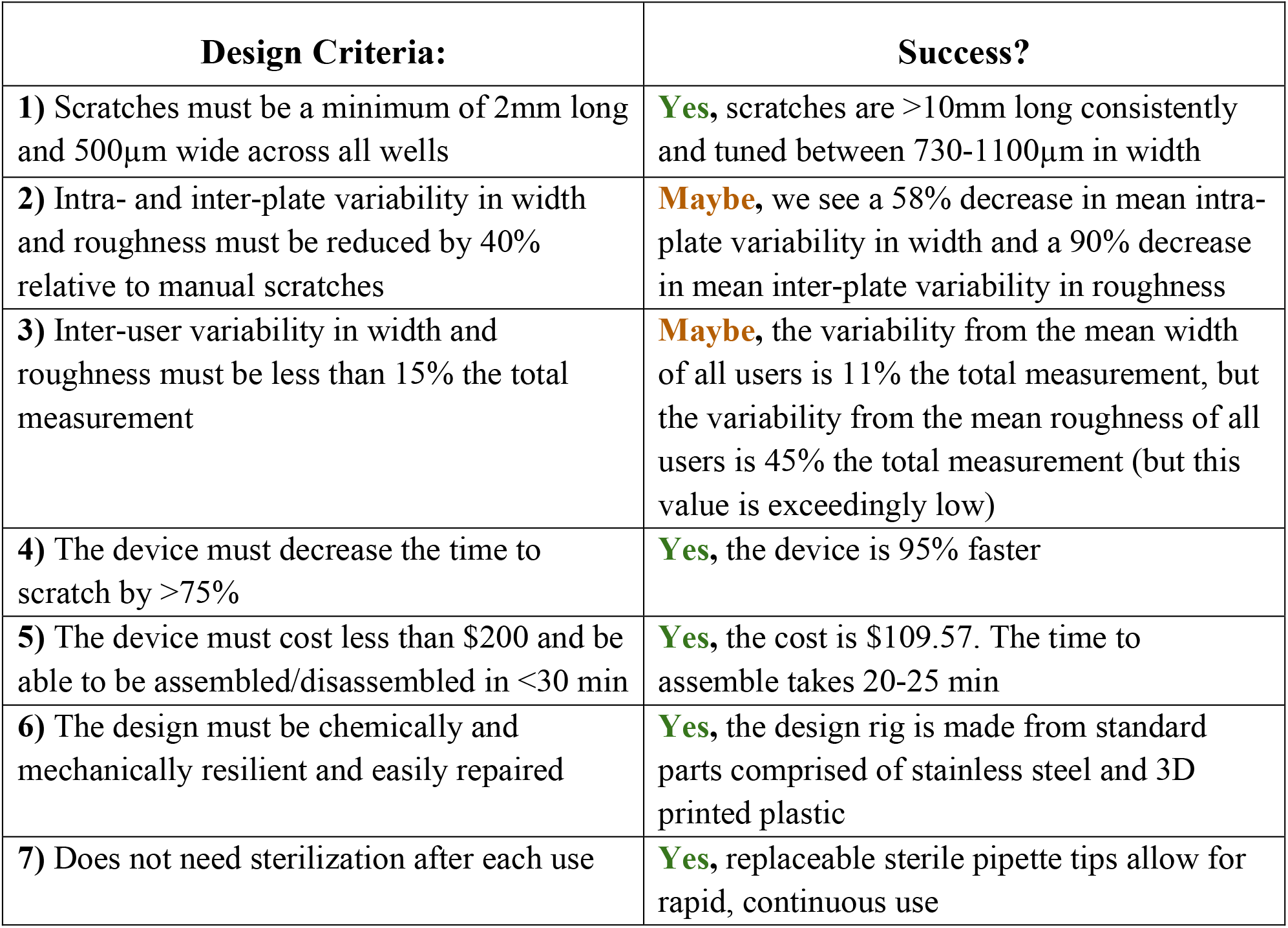
Evaluation of scratch assay device using design criteria

| **Design Criteria:** | **Success?** |
| --- | --- |
| **1)** Scratches must be a minimum of 2mm long and 500µm wide across all wells | **Yes**, scratches are >10mm long consistently and tuned between 730-1100µm in width |
| **2)** Intra- and inter-plate variability in width and roughness must be reduced by 40% relative to manual scratches | **Maybe**, we see a 58% decrease in mean intra-plate variability in width and a 90% decrease in mean inter-plate variability in roughness |
| **3)** Inter-user variability in width and roughness must be less than 15% the total measurement | **Maybe**, the variability from the mean width of all users is 11% the total measurement, but the variability from the mean roughness of all users is 45% the total measurement (but this value is exceedingly low) |
| **4)** The device must decrease the time to scratch by >75% | **Yes**, the device is 95% faster |
| **5)** The device must cost less than \$200 and be able to be assembled/disassembled in <30 min | **Yes**, the cost is \$109.57. The time to assemble takes 20-25 min |
| **6)** The design must be chemically and mechanically resilient and easily repaired | **Yes**, the design rig is made from standard parts comprised of stainless steel and 3D printed plastic |
| **7)** Does not need sterilization after each use | **Yes**, replaceable sterile pipette tips allow for rapid, continuous use |

Statistics: For **A, D**, and **G**, a Welch’s t-test (Gaussian distribution with unequal SD’s) was performed. For **H** an unpaired t-test (Gaussian distribution with equal SD’s) was performed.

Not significant (ns) and ***P< 0.0001 across all conditions. For **B, C, E**, and **F** a one-sample t-test was performed to compare the mean value against a threshold of 2500 µm^2^ (**B, C**) or 100 µm^2^ (**E, F**). Not significant (ns), *P < 0.03, **P < 0.008, and ***P ≤ 0.0003 across all conditions.

### 3.5 The scratch assay performed using the optimized scratch rig can detect differences in wound healing outcomes between unique cell treatments *in vitro*

Having optimized the Design 3 Scratch Rig and validated its utility against manual scratching, we wanted to assess the device under simulated *in vitro* use conditions. In the context of CNS injury, the ability of astrocytes to dedifferentiate into immature, neural progenitor-like cells, proliferate, and migrate into the site of injury ultimately influences the wound repair outcomes. Therefore, we wanted to test the ability of our device to reproducibly create scratches on plates of seeded neural progenitor cells (NPCs) and measure the response of the cells after injury when treated with media conditions that might alter their proliferative and migratory responses.

To do this, we seeded NPCs on 24-well plates and applied three different media conditions to the cells for 24 hours. By exposing NPCs to standard neural expansion media, conditioned with either 100ng/mL E/F (epithelial growth factor/fibroblast growth factor), 2% FBS (fetal bovine serum) with no added growth factors, or 50 E/Fng/mL + 2% FBS, we could create three different proliferative states in the NPCs. In our previous work, we have established that NPCs treated with E/F are highly proliferative and NPCs treated with FBS differentiate into quiescent astrocytes, becoming minimally proliferative. We have not previously assessed the effect of NPCs treated with both E/F and FBS concurrently in the context of scratch assay wound repair. Scratches were performed using the optimized scratch device, wound closure was then monitored via brightfield microscopy over a period of 54 hours with six separate imaging sessions conducted within that interval (**Fig. 5A**), and analyzed by using custom MATLAB code to find scratch width (**Fig. S8**).

**Figure 5:**
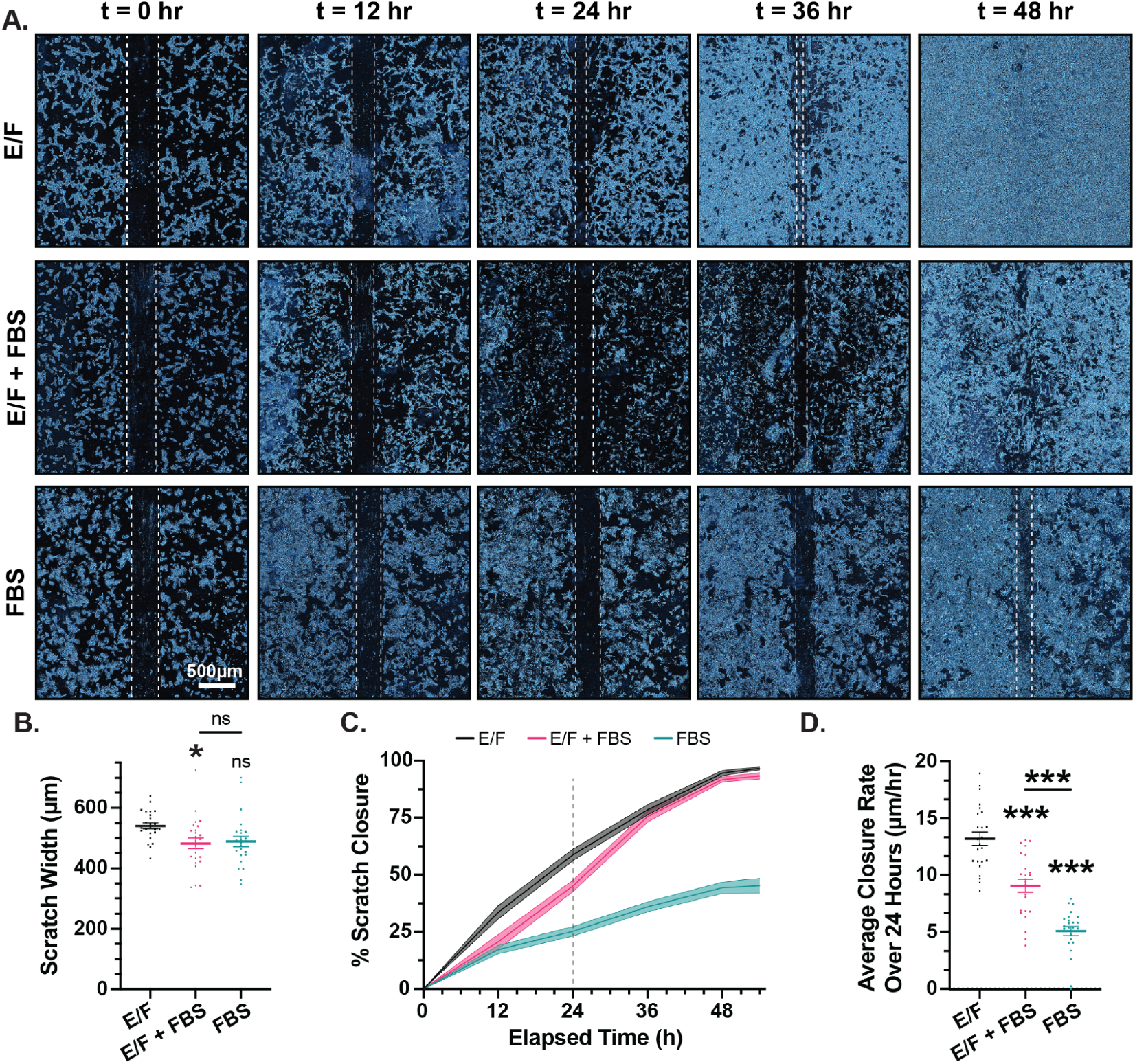
Scratch assay rig can be readily integrated into *in vitro* workflow and allows for detection of quantifiable differences between NPCs treated with E/F, E/F + FBS, and FBS. **A**. Timelapse images showing representative wound closure over time. Images were taken on a brightfield microscope, pseudocolored in FIJI, contrast enhanced using CLAHE filter, despeckled, and brightness/contrast adjusted for clarity. White dashed line highlights scratch boundaries. **B**. Plot showing the scratch width across the cell treatment conditions. Statistical comparisons to E/F or as designated. Not significant (ns) or *P< 0.03, one-way ANOVA with Tukey’s multiple comparison test. Graph shows mean ± s.e.m. with individual data points showing n = 24 wells. **C**. Trace showing % Scratch Closure over time, dashed line marks 24 hour time point. Graph shows mean ± s.e.m. with s.e.m. represented as a light shaded area. n=24 per group. **D**. Plot showing the effect of cell treatment conditions on scratch closure rate within the first 24 hours. Statistical comparisons to E/F or as designated. ***P< 0.0001, one-way ANOVA with Tukey’s multiple comparison test. Graph shows mean ± s.e.m. with individual data points showing n = 24 wells.

Immediately after scratching, the wounds had an average width of 483-541 µm across the three conditions (**Fig. 5A-B**). Interestingly, these scratches were significantly thinner than those created in the dry erase assay, suggesting that the dry erase assay may not fully replicate the physical characteristics of *in vitro* wounds. By tracking the average width of the scratch over time and normalizing it to the initial scratch width, we can evaluate the percentage scratch closure. Scratches made on cells treated with E/F and E/F + FBS closed quickly, with nearly complete cellular repopulation of the scratched region after 48 hrs (**Fig. 5C**). Meanwhile, the lowly proliferative, FBS-treated NPCs could not close the wound fully over the course of 54hrs (**Fig. 5C**). By fitting a simple linear model to the first 24hrs, we determine the rate of wound closure in µm/hr (**Fig. 5D**). NPCs treated with E/F had the highest rate of closure at 13.2 µm/hr, which was significantly greater than E/F + FBS (9.1 µm/hr), and FBS (5.1 µm/hr) (**Fig. 5D**). Collectively, this data demonstrates that the Scratch Assay Rig is compatible with *in vitro* models of wound healing and provides reproducible scratches with sufficiently low variability such that differences in wound repair functions of NPCs could be readily identified.

## 4. Discussion

Mechanically-driven central nervous system (CNS) injuries, such as traumatic brain injury (TBI) and spinal cord injury (SCI), cause widespread neural tissue damage that disrupts homeostatic function [1–4]. These injuries trigger a complex, coordinated multicellular response involving immune cells and the endogenous glial cells [1,4]. The success of this response in restoring neural function is dependent on the interplay between pro-fibrotic, immune-based repair and endogenous, glia-mediated repair mechanisms [5,6]. However, endogenous repair mechanisms are often insufficient to fully restore neural function, underscoring the need to dissect glial repair processes to develop mechanistic insights and identify therapeutic strategies to enhance recovery outcomes.

Scratch assays offer an elegant approach to assess mechanisms of wound repair by isolating specific cell types and allowing for precise regulation of the cellular microenvironment [9]. These assays also allow for rapid testing of treatment strategies to enhance glial repair by augmenting cellular proliferation and migration. Nevertheless, traditional scratch assays suffer from low throughput and poor reproducibility, which stems from varied scratch width and high scratch tortuosity both between wells and users. Here, we report the fabrication of an open-source scratch assay device that can be readily scaled to scratch 6-well plates to 96-well plates. This scratch assay device can be fabricated for less than $200 and enhances the scratch speed by 95%, while the standard deviation of tortuosity and roughness decreased by 90% and 47%, respectively (relative to manual scratches). This device has broad applications for wound healing studies, and future work should examine: (1) refining the open-source design by exploring alternatives to Amazon-sourced components; (2) developing methods to better tune scratch width; and (3) screening molecules to enhance glial proliferation and migration capacity.

We are enthusiastic about the design of our scratch assay device. By combining readily available components from Amazon and McMaster-Carr with a few custom, 3D-printed parts, we have created a device that can be fabricated for $109.57 without requiring any specialized expertise. While this product represents a significant improvement on current scratch devices, future iterations could be enhanced by: (1) converting all components to 3D-printed parts or (2) sourcing all components from a single, larger vendor, such as McMaster-Carr. Presently, the actuating scratch plate is sourced from Amazon, which offers the benefits of low cost and rapid delivery, but lacks a reliable, long-term supply chain. To ensure sustainable fabrication of our scratch assay device, it would be ideal to design the base to be 3D-printed. This would both serve to secure the supply of actuating bases, but would also improve device reproducibility by producing identical bases for each build and eliminating the need for machining. A second approach to address sourcing concerns would be to establish a design for this device using all components from a single industrial vendor, like McMaster-Carr. While this does not provide immunity from future changes in the production of those components, industrial distributors like McMaster-Carr have the ability to source alternate parts meeting customer specifications, allowing for the rapid substitution of comparable components.

A second avenue to investigate with this device is the development of more robust methods to tune scratch width. In this study, we demonstrated that adjusting the angle of the pipette tips can modulate scratch width between 681 µm and 864 µm, with minimal impact on inter-or intra-plate variability. However, this ∼180 µm of tunability is considerably limited. Alternatively, altering the pipette tip geometry by shortening the tip length allowed for scratches ranging from 730 µm to 1100 µm in width, but this approach increased inter-and intra-plate variability, reducing the reproducibility of these wider scratches. To overcome these limitations, future work could explore redesigning the tip holders for differently sized orifice tips (such as p10 or p1000 tips). This modification would enable significantly narrower or wider scratches, while also leveraging the modular nature of this rig design. Another potential strategy would be designing a pipette tip guillotine to produce consistent orifice widths and geometries. We hypothesize that the increased variability observed with wider bore tips arises from combined differences in orifice diameter, pipette tip length, and the relative angle of the cut. Therefore, ensuring consistency in these variables through the design of a secondary pipette tip trimming device should reduce scratch width variability and improve reproducibility for these larger scratches.

Validation of this assay with NPCs under varied media conditions demonstrates the scratch device’s ability to create reproducible outcomes that can discriminate between rapid and limited wound healing cell capacities. Having characterized the efficacy of this assay, we are excited to test various metabolically active molecules [22–24] and growth factors [25–27] to assess their impact on astrocyte proliferation, migration, and wound repair capabilities. Using our newly developed scratch assay rig will ultimately enable hypothesis-driven, fundamental biology studies within a high-throughput, cost-effective experimental framework, facilitating rapid mechanistic insights and subsequent translation to *in vivo* models.

## 5. Conclusion

In conclusion, this work describes the design and implementation of a low cost, open source, scratch assay device. Our results show that this scratch assay device can be fabricated for $110 and enhances scratch reproducibility, while increasing throughput by 95%. This work has promise for use both in examining the effects of metabolically active and proliferation-enhancing molecules on wound healing in the CNS and more generally for use in other classic wound healing or migration-based experiments such as in the context of diabetes [28,29] and cancer [30,31].

## Abbreviations

CNS: Central Nervous System
BBB: Blood Brain Barrier
NPCs: Neural Progenitor Cells
TBI: Traumatic Brain Injury
SCI: Spinal Cord Injury
CAD: Computer-Aided Design
PLA: Polylactic Acid
NE: Neural Expansion
FBS: Fetal Bovine Serum
FDM: Fused Deposition Modeling
TC: Tissue Culture
FOV: Field of View
EGF: Epidermal Growth Factor
FGF: Fibroblast Growth Factor
PBS: Phosphate Buffered Saline

## Acknowledgements

The authors are grateful to Eliott Dinfotan for the generous use of his Creality printer and freely given printing advice. We would also like to thank Alexander Rocca for additional 3D printing guidance, and Isaac Herrick and Sebastian Alfreds for their help in testing user effect on the scratch assay device. This research was also supported by the Micro and Nano Imaging (MNI) core in the Biomedical Engineering Core Facilities at Boston University and the Singh Imagineering (SI) Lab in the College of Engineering at Boston University. Finally, we would like to thank the many funding sources that contributed to this work including Boston University Start-Up Funds, Boston University’s Distinguished Summer Research Fellowship (DSRF), Boston University’s Undergraduate Research Opportunities Program (UROP), and Ruth L. Kirschstein Predoctoral Individual National Research Service Award, NIH NINDS (F31NS135944 to E.M.D), Maximizing Investigators’ Research Award (MIRA), NIH NIGMS (R35GM154942 to T.M.O.) (the content in this manuscript is solely the responsibility of the authors and does not reflect the official views of the NIH).

## Author Contributions

**CRediT (Contributor Roles Taxonomy) roles:**

- **Conceptualization -** J.E.L. + E.M.D. + T.M.O.
- **Data curation -** E.M.D. + T.M.O.
- **Formal analysis -** P.K. + E.M.D. + T.M.O.
- **Funding acquisition -** J.E.L. + P.K. + T.M.O.
- **Investigation -** J.E.L. + P.K. + E.M.D. + K.Y. + T.M.O.
- **Methodology -** J.E.L. + P.K. + E.M.D. + K.Y. + T.M.O.
- **Project administration -** E.M.D. + T.M.O.
- **Resources -** J.E.L. + P.K. + T.M.O.
- **Software -** P.K.
- **Supervision -** E.M.D. + T.M.O.
- **Validation -** P.K. + E.M.D. + T.M.O.
- **Visualization -** P.K. + E.M.D. + T.M.O.
- **Writing – original draft -** J.E.L. + P.K. + E.M.D. + T.M.O.
- **Writing – review and editing -** J.E.L. + P.K. + E.M.D. + T.M.O.

## References

[1] O’Shea, T.M. et al. (2017). Cell biology of spinal cord injury and repair. The Journal of Clinical Investigation. 10.1172/JCI90608.

[2] Venkatesh, K. et al. (2019). Spinal cord injury: pathophysiology, treatment strategies, associated challenges, and future implications. Cell and Tissue Research. 10.1007/s00441-019-03039-1.

[3] Ng, S.Y. and Lee, A.Y.W. (2019). Traumatic Brain Injuries: Pathophysiology and Potential Therapeutic Targets. Frontiers in Cellular Neuroscience. 10.3389/fncel.2019.00528.

[4] Mira, R.G. et al. (2021). Traumatic Brain Injury: Mechanisms of Glial Response. Frontiers in Physiology. 10.3389/fphys.2021.740939.

[5] Bradbury, E.J. and Burnside, E.R. (2019). Moving beyond the glial scar for spinal cord repair. Nature Communications. 10.1038/s41467-019-11707-7.

[6] Dias, D.O. and Göritz, C. (2018). Fibrotic scarring following lesions to the central nervous system. Matrix Biology. 10.1016/j.matbio.2018.02.009.

[7] Li, Y. et al. (2020). Microglia-organized scar-free spinal cord repair in neonatal mice. Nature. 10.1038/s41586-020-2795-6.

[8] Skinnider, M.A. et al. (2024). Single-cell and spatial atlases of spinal cord injury in the Tabulae Paralytica. Nature. 10.1038/s41586-024-07504-y.

[9] Liang, C.-C. et al. (2007). In vitro scratch assay: a convenient and inexpensive method for analysis of cell migration in vitro. Nature Protocols. 10.1038/nprot.2007.30.

[10] Stamm, A. et al. (2016). In vitro wound healing assays – state of the art. BioNanoMaterials. 10.1515/bnm-2016-0002.

[11] Cappiello, F. et al. (2018). A Novel In Vitro Wound Healing Assay to Evaluate Cell Migration. Journal of Visualized Experiments : JoVE. 10.3791/56825.

[12] Planz, V. et al. (2018). Establishment of a cell-based wound healing assay for bio-relevant testing of wound therapeutics. Journal of Pharmacological and Toxicological Methods. 10.1016/j.vascn.2017.10.003.

[13] Lan, R. et al. (2010). A Novel Wounding Device Suitable for Quantitative Biochemical Analysis of Wound Healing and Regeneration of Cultured Epithelium. Wound repair and regeneration : official publication of the Wound Healing Society [and] the European Tissue Repair Society. 10.1111/j.1524-475X.2010.00576.x.

[14] Lee, J. et al. (2010). Stamp Wound Assay for Studying Coupled Cell Migration and Cell Debris Clearance. Langmuir. 10.1021/la103542y.

[15] Zordan, M.D. et al. (2011). A High Throughput, Interactive Imaging, Bright-Field Wound Healing Assay. Cytometry. 10.1002/cyto.a.21029.

[16] Yue, P.Y.K. et al. (2010). A Simplified Method for Quantifying Cell Migration/Wound Healing in 96-Well Plates. Journal of Biomolecular Screening. 10.1177/1087057110361772.

[17] Yarrow, J.C. et al. (2004). A high-throughput cell migration assay using scratch wound healing, a comparison of image-based readout methods. BMC Biotechnology. 10.1186/1472-6750-4-21.

[18] Chen, S. et al. (2023). An effective device to enable consistent scratches for in vitro scratch assays. BMC Biotechnology. 10.1186/s12896-023-00806-5.

[19] Grimmig, R. et al. (2019). Development and Evaluation of a Prototype Scratch Apparatus for Wound Assays Adjustable to Different Forces and Substrates. Applied Sciences. 10.3390/app9204414.

[20] Lin, Y. et al. (2024). A programmable, open-source robot that scratches cultured tissues to investigate cell migration, healing, and tissue sculpting. Cell Reports Methods. 10.1016/j.crmeth.2024.100915.

[21] O’Shea, T.M. et al. (2022). Lesion environments direct transplanted neural progenitors towards a wound repair astroglial phenotype in mice. Nature Communications. 10.1038/s41467-022-33382-x.

[22] Zhang, Y. et al. (2023). Astrocyte metabolism and signaling pathways in the CNS. Frontiers in Neuroscience. 10.3389/fnins.2023.1217451.

[23] Xiong, X.-Y. et al. (2022). Metabolic changes favor the activity and heterogeneity of reactive astrocytes. Trends in Endocrinology & Metabolism. 10.1016/j.tem.2022.03.001.

[24] Theparambil, S.M. et al. (2024). Adenosine signalling to astrocytes coordinates brain metabolism and function. Nature. 10.1038/s41586-024-07611-w.

[25] Savchenko, E. et al. (2019). FGF family members differentially regulate maturation and proliferation of stem cell-derived astrocytes. Scientific Reports. 10.1038/s41598-019-46110-1.

[26] Prah, J. et al. (2019). A novel serum free primary astrocyte culture method that mimic quiescent astrocyte phenotype. Journal of Neuroscience Methods. 10.1016/j.jneumeth.2019.03.013.

[27] Pöyhönen, S. et al. (2019). Effects of Neurotrophic Factors in Glial Cells in the Central Nervous System: Expression and Properties in Neurodegeneration and Injury. Frontiers in Physiology. 10.3389/fphys.2019.00486.

[28] Soheilifar, M.H. et al. (2024). In vitro and in vivo evaluation of the diabetic wound healing properties of Saffron (Crocus Sativus L.) petals. Scientific Reports. 10.1038/s41598-024-70010-8.

[29] Li, L. et al. (2019). High Glucose Suppresses Keratinocyte Migration Through the Inhibition of p38 MAPK/Autophagy Pathway. Frontiers in Physiology. 10.3389/fphys.2019.00024.

[30] Wang, Y. et al. (2025). Mechanisms of C5ORF13 in promoting epithelial-mesenchymal transition in hepatocellular carcinoma via eIF6 and the intervention of valproic acid. Pathology - Research and Practice. 10.1016/j.prp.2025.156211.

[31] Zhang, J. et al. (2025). The peptide ENSG00000232259-ORF inhibits glioma cell proliferation and migration by promoting autophagy. Brain Research. 10.1016/j.brainres.2025.149942.

