## Appendix A for "In Vitro Wound Simulation: A High-Throughput Device for Scratch Assays"

**Appendix A. Supplementary Data**

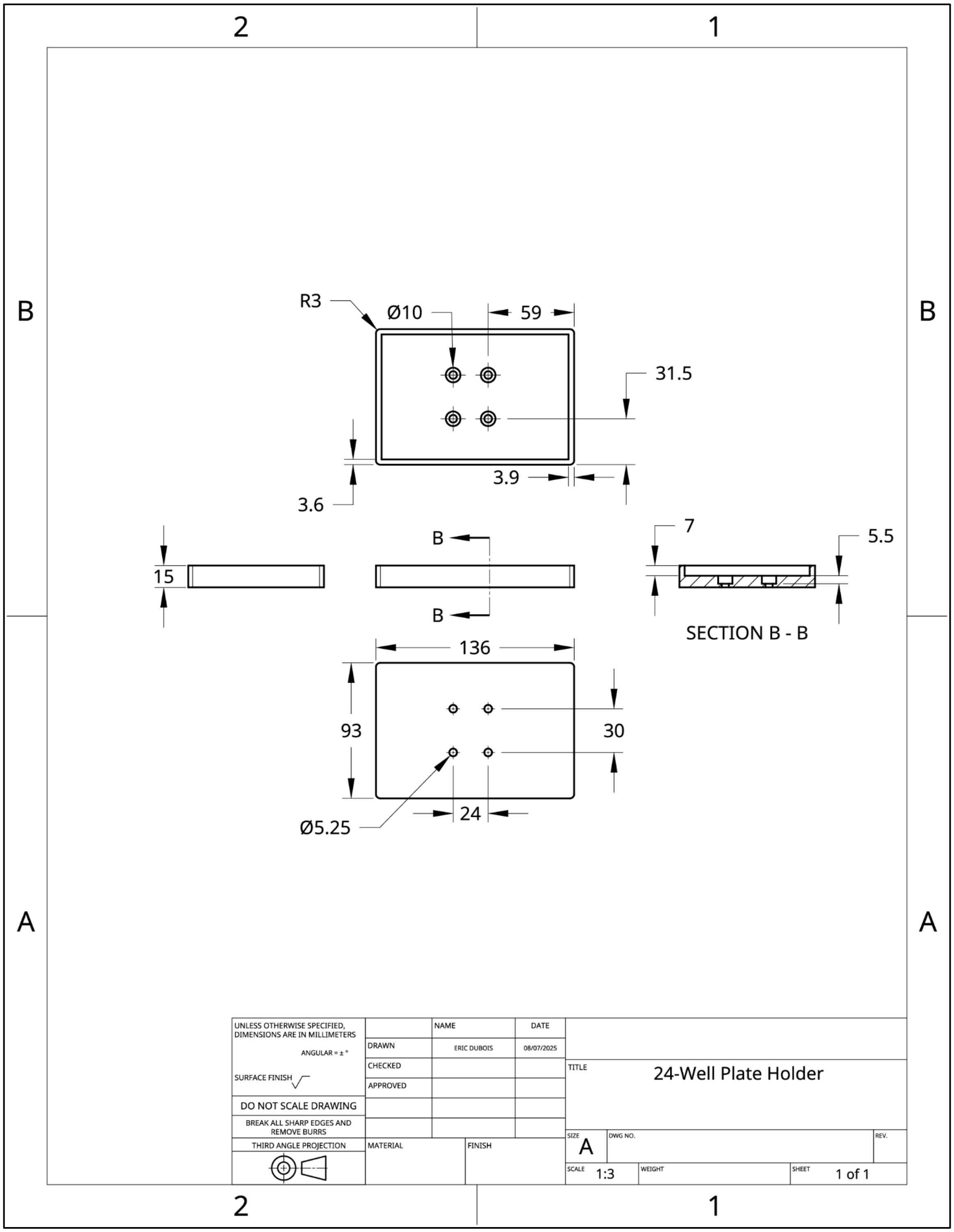


**Figure S1: Drawing for 24-Well Plate Holder.** Drawing uses ANSI part drawing template from Onshape. All units are in mm unless otherwise noted.

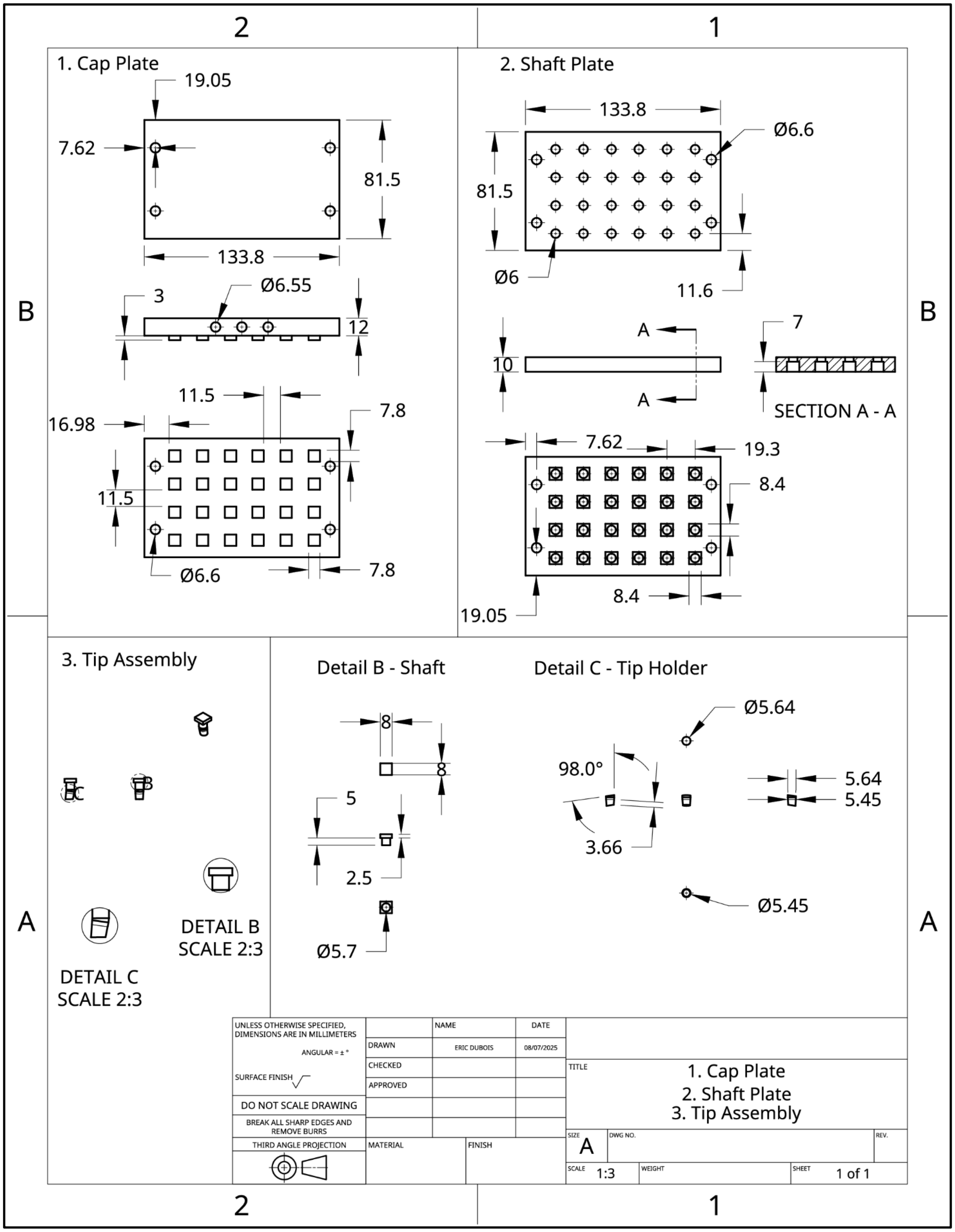


**Figure S2: Drawing for Design 3.** All three individual components are on this drawing, including the: (1) Cap Plate, (2) Shaft Plate, and (3) Tip Assembly. Drawing uses ANSI part drawing template from Onshape. All units are in mm unless otherwise noted.

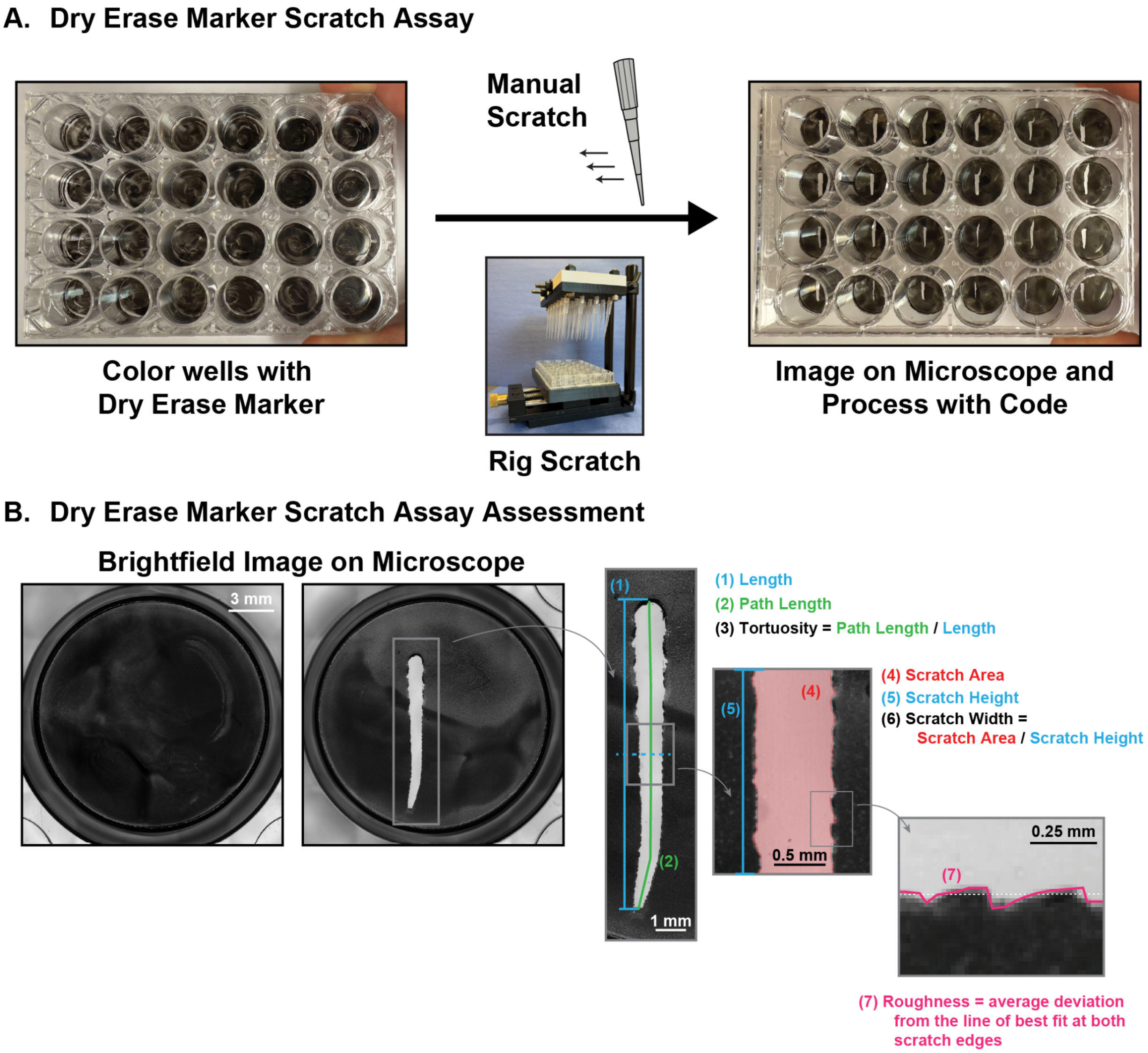


**Figure S3: Dry Erase Marker Scratch Assay Design and Testing. A.** Images showing the 24-well plate after coating with dry erase marker, prior to scratching with either the manual scratch technique or the rig, and after scratching. **B.** Brightfield images on microscope before and after scratching. Scratch image has been annotated to highlight the key features extracted and reported in this work, including: (1) length, (3) tortuosity, (6) scratch width, and (7) roughness.

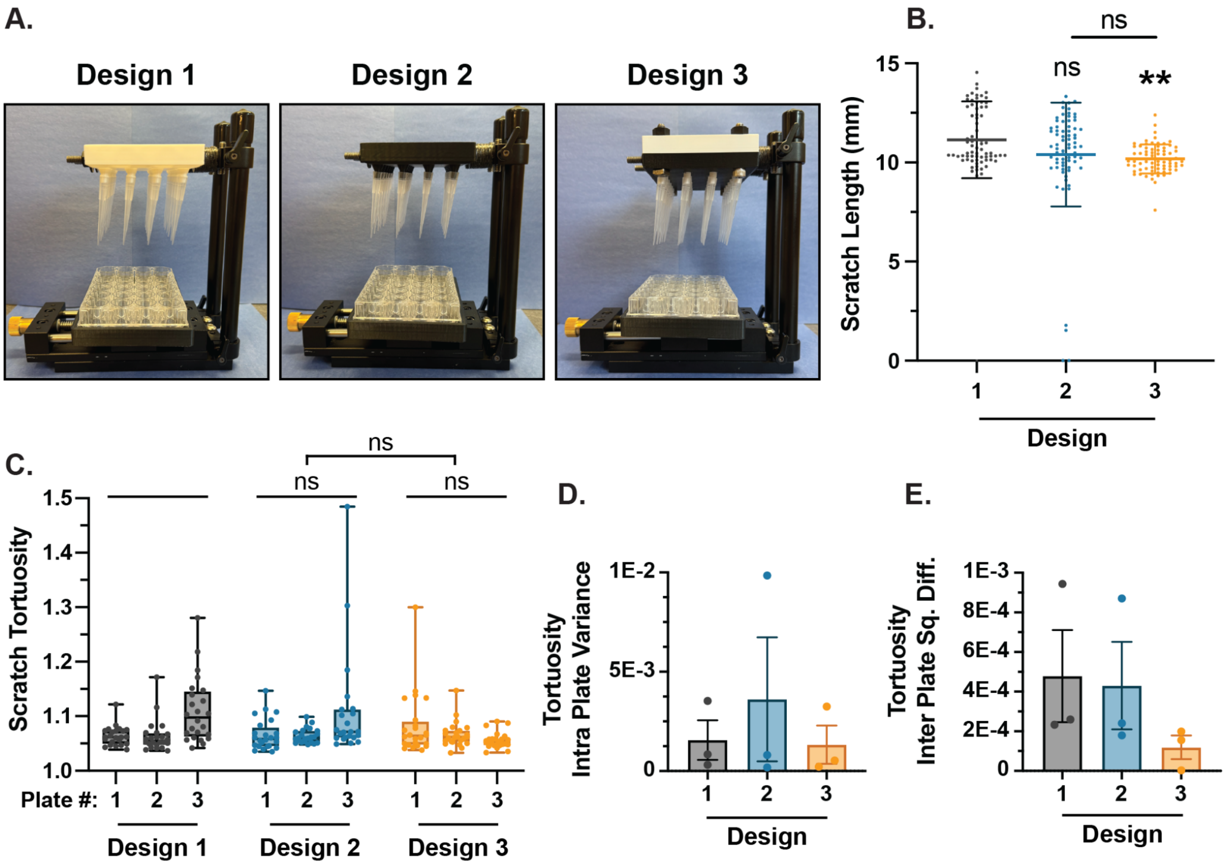


**Figure S4: Effect of Design on scratch length and tortuosity. A.** Images showing the completed Design 1-3. **B.** Graph showing the effect of design iteration on scratch length. Statistical comparisons to Design 1 or as designated. Graph shows mean ± SD with individual data points showing n = 72 wells from three 24-well plates. **C.** Box and whisker plot showing the effect of design on scratch tortuosity. Statistical comparisons to Design 1 or as designated. Box shows mean and interquartile range (IQR), whiskers show full data range with individual data points showing n = 24 wells per plate, except for Design 1 Plate 1, and Design 2 Plate 2-3 (n = 23). **D.** Bar graph showing the effect of design iteration on the intra-plate variance for scratch tortuosity. Graph shows mean ± s.e.m. with individual data points showing n = 3 plates. **E.** Bar graph showing the effect of design iteration on the inter-plate squared differences (Sq. Diff.) for scratch tortuosity. Graph shows mean ± s.e.m. with individual data points showing n = 3 plates.

Statistics: For **B** a one-way ANOVA (Gaussian distribution with equal SD's) was performed followed by a Tukey's multiple comparison test. For **C** a Brown-Forsythe and Welch ANOVA test (Gaussian distribution with unequal SD's) was performed followed by a Dunnett's T3 multiple comparisons test. Not significant (ns) and $^{**}P < 0.008$ across all conditions.

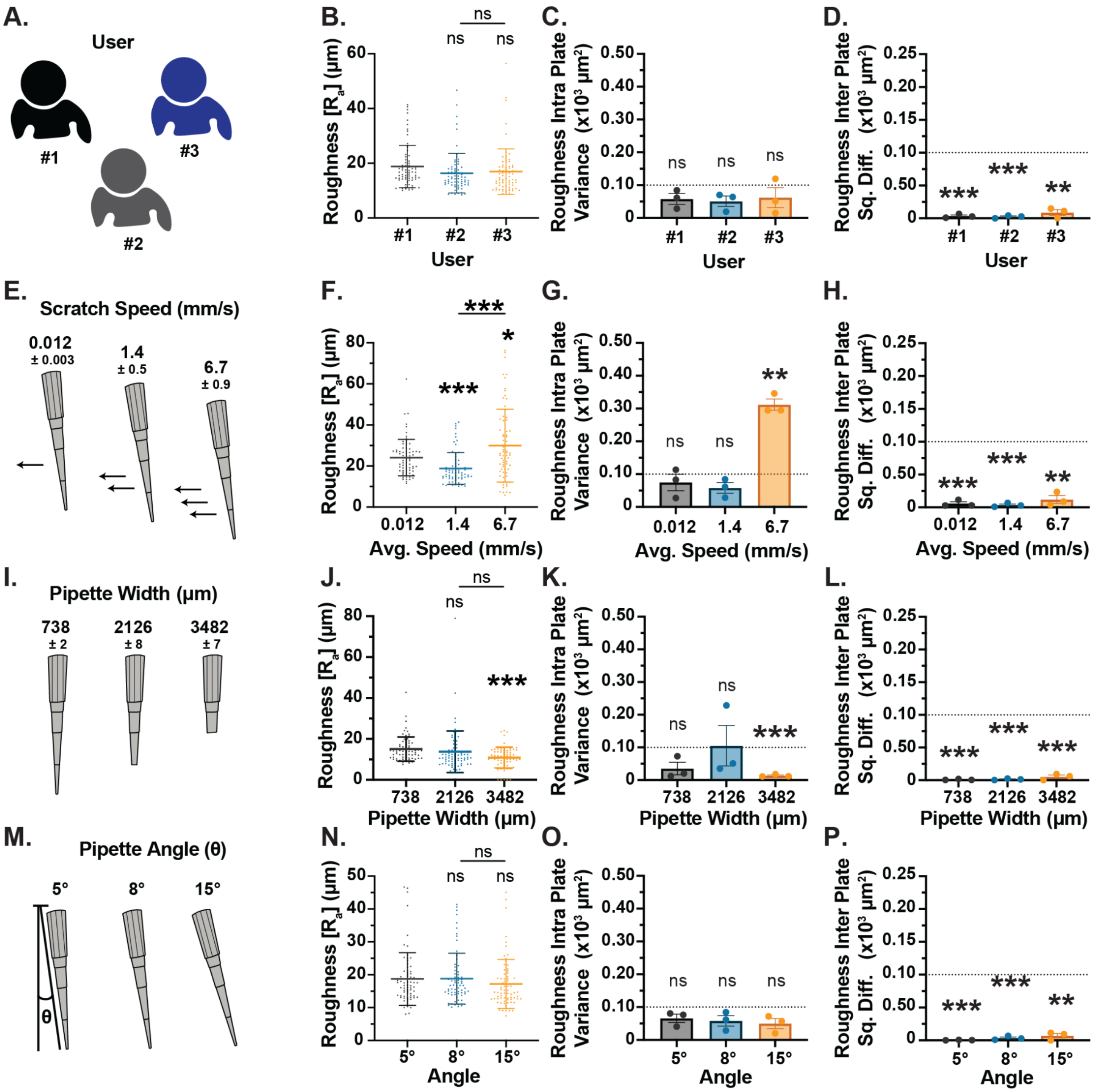


**Figure S5: Design 3 scratch rig creates controlled scratches that can be tuned using simple tip parameters. A.** Schematic showing tested variable: User. **B.** Graph showing the effect of user on scratch roughness. Statistical comparisons to User #1 or as designated. Graph shows mean ± SD with individual data points showing n = 72 wells from three 24-well plates. **C.** Bar graph showing the effect of user on the intra-plate variance for scratch roughness. Not significant (ns) across all conditions. Graph shows mean ± s.e.m. with individual data points showing n = 3 plates. **D.** Bar graph showing the effect of user on the inter-plate squared differences (Sq. Diff.) for scratch roughness. Not significant (ns) across all conditions. Graph shows mean ± s.e.m. with individual data points showing n = 3 plates. **E.** Schematic showing tested variable: Scratch Speed. Reported values are the mean ± the 95% confidence interval (CI). **F.** Graph showing the effect of scratch speed on scratch roughness. Statistical comparisons to 0.012 (mm/s) or as designated. Graph shows mean ± SD with individual data points showing n = 72 wells from three 24-well plates. **G.** Bar graph showing the effect of scratch speed on the intra-plate variance for scratch roughness. Not significant (ns) and *P < 0.05 across all conditions. Graph shows mean ± s.e.m. with individual data points showing n = 3 plates. **H.** Bar graph showing the effect of scratch speed on the inter-plate squared differences (Sq. Diff.) for scratch roughness. Not significant (ns) across all conditions. Graph shows mean ± s.e.m. with individual data points showing n = 3 plates. **I.** Schematic showing tested variable: Pipette Width. Reported values are the mean ± the 95% confidence interval (CI). **J.** Graph showing the effect of pipette width on scratch roughness. Statistical comparisons to narrowest width or as designated. Graph shows mean ± SD with individual data points showing n = 72 wells from three 24-well plates. **K.** Bar graph showing the effect of pipette width on the intra-plate variance for scratch roughness. Not significant (ns) across all conditions. Graph shows mean ± s.e.m. with individual data points showing n = 3 plates. **L.** Bar graph showing the effect of pipette width on the inter-plate squared differences (Sq. Diff.) for scratch roughness. Not significant (ns) across all conditions. Graph shows mean ± s.e.m. with individual data points showing n = 3 plates. **M.** Schematic showing tested variable: Pipette Angle. **N.** Graph showing the effect of pipette angle on scratch roughness. Statistical comparisons to 5° or as designated. Graph shows mean ± SD with individual data points showing n = 72 wells from three 24-well plates. **O.** Bar graph showing the effect of pipette angle on the intra-plate variance for scratch roughness. Not significant (ns) across all conditions. Graph shows mean ± s.e.m. with individual data points showing n = 3 plates. **P.** Bar graph showing the effect of pipette angle on the inter-plate squared differences (Sq. Diff.) for scratch roughness. Not significant (ns) across all conditions. Graph shows mean ± s.e.m. with individual data points showing n = 3 plates.

Statistics: For **B** and **N** a one-way ANOVA (Gaussian distribution with equal SD's) was performed followed by a Tukey's multiple comparison test. For **F** and **J** a Brown-Forsythe and Welch ANOVA test (Gaussian distribution with unequal SD's) was performed followed by a Dunnett's T3 multiple comparisons test. Not significant (ns), *P< 0.05, and ***P< 0.0008 across all conditions. For **C, D, G, H, K, L, O** and **P** a one-sample t-test was performed to compare the mean value against a threshold of 100 $\mu m^2$. Not significant (ns), **P < 0.007 , and ***P ≤ 0.0009 across all conditions.

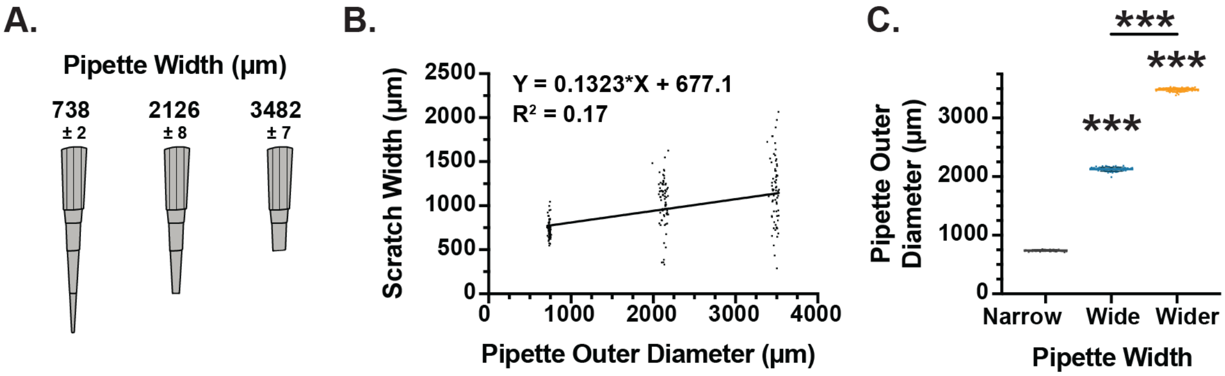


**Figure S6: Pipette widths are minimally variable after trimming. A.** Schematic showing tested variable: Pipette Width. Reported values are the mean ± the 95% confidence interval (CI). **B.** Plot showing the effect of pipette outer diameter on scratch width. Equation gives the line of best fit for a simple linear regression and $R^2$ value is from a Pearson's Correlation Test. n = 216 total data points (72 wells / pipette width). **C.** Plot showing the consistency in pipette outer diameters. Outer diameters are created by trimming pipette tips to create narrow (untrimmed), wide (small trim), and wider (largest trim). Statistical comparisons to Narrow or as designated. ***P≤ 0.0001, Brown-Forsythe and Welch ANOVA test (Gaussian distribution with unequal SD's) followed by a Dunnett's T3 multiple comparisons test. Graph shows mean ± SD with individual data points showing n = 72 wells per condition.

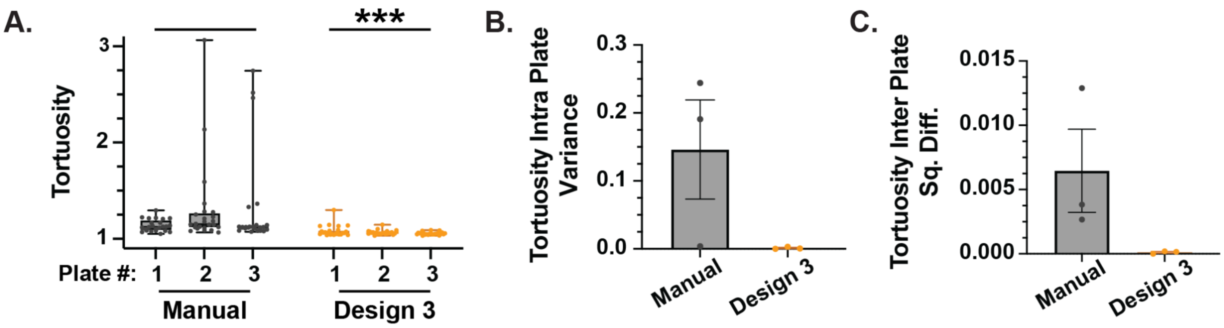


**Figure S7: Design 3 Scratch Rig minimizes tortuosity when compared to a manual scratch. A.** Box and whisker plot showing the effect of scratch rig implementation on scratch tortuosity. ***P≤ 0.0001 between the two conditions, Welch's t-test. Box shows mean and interquartile range (IQR), whiskers show full data range with individual data points showing n = 24 wells per plate. **B.** Bar graph showing the effect of scratch rig implementation on the intra-plate variance for scratch tortuosity. Graph shows mean ± s.e.m. with individual data points showing n = 3 plates. **C.** Bar graph showing the effect of scratch rig implementation on the inter-plate squared differences (Sq. Diff.) for scratch tortuosity. Graph shows mean ± s.e.m. with individual data points showing n = 3 plates.

**Table S1**: Cost analysis of building the scratch assay device

| **Index** | **Item Name** | **Source** | **Product Number** | **Bulk Cost** | **Units in Bulk Order** | **Mass (g)** | **Units Needed for Rig** | **Cost for Device** |
|---|---|---|---|---|---|---|---|---|
| **1** | Base Plate | Amazon | SmallRig 1092 | $27.99 | 1 | | 1 | $27.99 |
| **2** | Actuating Plate | Amazon | B09HQGTDFT | $39.79 | 1 | - | 1 | $39.79 |
| **3** | Rod Clamp | Amazon | SmallRig 2061 | $14.99 | 2 | - | 2 | $14.99 |
| **4** | Rods | Amazon | B082LSTSN2 | $12.59 | 2 | - | 2 | $12.59 |
| **5** | Rubber Caps | McMaster-Carr | 6448K32 | $7.76 | 10 | - | 2 | $1.55 |
| **6** | ¼”-20 Threaded Rods, 5” | McMaster-Carr | 91565A562 | $8.64 | 10 | - | 2 | $1.73 |
| **7** | ¼” Nylon Washers | McMaster-Carr | 90295A455 | $6.55 | 50 | - | 24 | $3.14 |
| **8** | ¼”-20 Nylon Nuts | McMaster-Carr | 94812A700 | $13.24 | 100 | - | 6 | $0.79 |
| **9** | ¼”-20, 3/8" long Set Screws | McMaster-Carr | 92311A535 | $10.14 | 100 | - | 4 | $0.41 |
| **10** | M5x0.8mm Screws, 12mm length | McMaster-Carr | **91292A125** | $12.62 | 100 | - | 4 | $0.50 |
| **11** | ¼”-20 Screws, 1 1/4” length | McMaster-Carr | 92196A544 | $19.77 | 25 | - | 4 | $3.16 |
| **12** | ¼”-20 Screws, 1/2” length | McMaster-Carr | 92196A537 | $20.71 | 50 | - | 2 | $0.83 |
| **13** | Foam earplugs | Amazon | 70356AB | $8.90 | 100 | - | 2 | $0.18 |
| **14** | Creality 1.75mm PLA filament | Amazon | 3301010337 | $23.99 | 2kg | - | 1 | - |
| **15** | *Well Plate Holder* | - | - | - | - | 50.1 | 1 | $0.60 |
| **16** | *Printed Shim Set* | - | - | - | - | 1.3 | 1 | $0.02 |
| **17** | *Shaft Plate* | - | - | - | - | 44.6 | 1 | $0.53 |
| **18** | *Cap Plate* | - | - | - | - | 53.6 | 1 | $0.64 |
| **19** | *Pipette Tip Holders* | - | - | - | - | 0.4 | 24 | $0.12 |

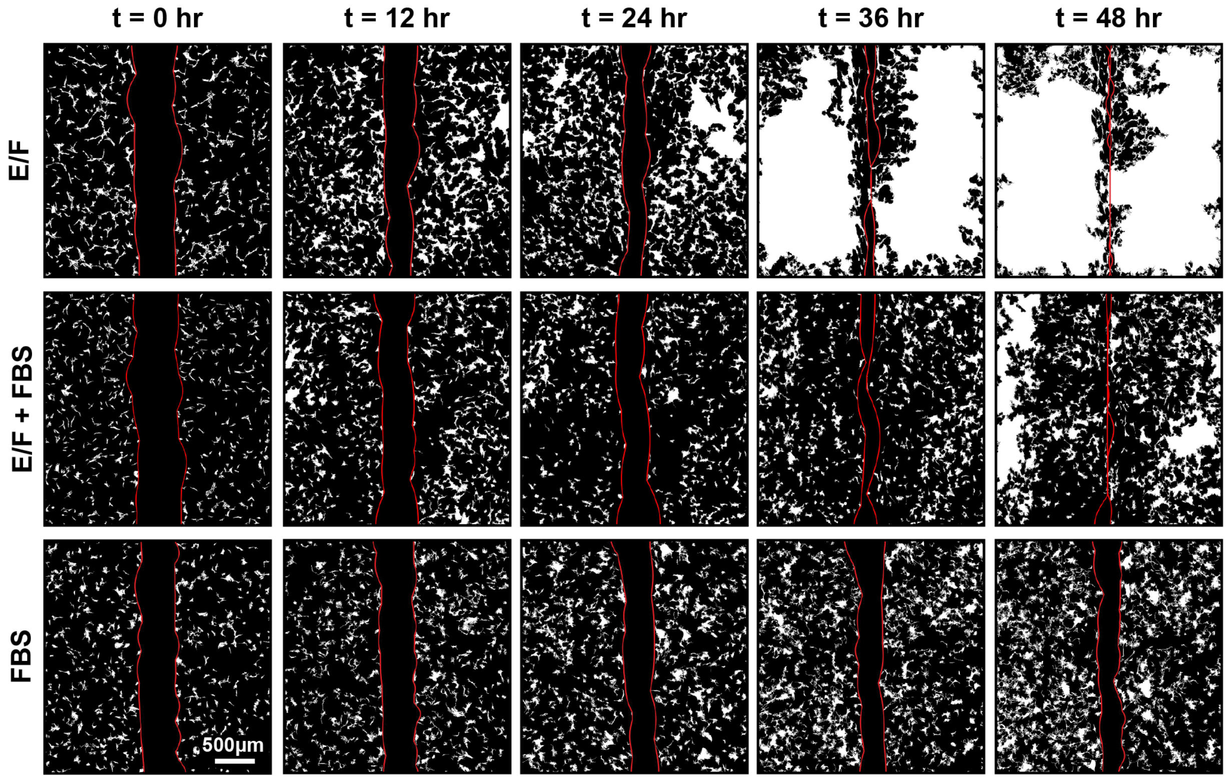


**Figure S8: Overview images of representative scratches for NPCs treated with E/F, E/F + FBS, and FBS.** Brightfield microscopy images that have been processed by a top-hat filter of 15 pixels and Li's threshold to yield a binarized image (white = detected cells). Red line demarcates the detected scratch, which was automatically determined by tracing cells and iteratively smoothing the produced curves in MATLAB.
